# Meta-analysis of the uncultured gut microbiome across 11,115 global metagenomes reveals a new candidate biomarker of health

**DOI:** 10.1101/2025.09.09.675081

**Authors:** Ana C. da Silva, Jacob Lapkin, Qi Yin, Efrat Muller, Alexandre Almeida

## Abstract

The human gut microbiome plays an important biological role in host health, yet over 60% of gut species remain uncultured and hence inaccessible to experimental manipulation. Here we analysed 11,115 human gut metagenomes from 39 countries, 13 noncommunicable diseases and healthy individuals to understand the clinical relevance of the uncultured microbiome worldwide. We identified 317 uncultured species linked to distinct health states, with most microbiome signatures being disease specific. Uncultured bacteria were more abundant in healthy controls, with the genus CAG-170 predicted as the strongest biomarker of health. The genetic diversity and abundance of CAG-170 negatively correlated with gut microbiome imbalance over time, and ecological modelling identified it as the top candidate keystone genus among healthy populations globally. Functional prediction analysis showed CAG-170 species have greater capabilities for vitamin B12 biosynthesis but lack key genes involved in arginine production, offering new biological insights into this elusive genus. Our findings shed light on the underexplored role of uncultured gut species in health and disease.

## Main

The human gut microbiome is a complex microbial ecosystem implicated in a variety of functions involved in host health, including nutrient metabolism, immune modulation, and colonization resistance against enteric pathogens^1,2^. Bacterial cultivation of gut microbiome species has been an essential tool for studying the role of the human microbiome in health, as it enables the experimental manipulation and testing of bacterial isolates under controlled laboratory conditions^3^. However, due to the complexity of the gut microbiome and the inherent challenges of culturing many of its constituent species, culture-independent methods, namely 16S rRNA gene sequencing and shotgun metagenomics, have been utilized to characterize microbiomes across different populations and health states^4,5^.

A large body of research applying 16S rRNA gene sequencing to microbiome data has linked specific microbial signatures to diseases such as inflammatory bowel disease, colorectal cancer, obesity, among others^6^. Although 16S rRNA genotyping has historically been the most widely used technique for microbiome analysis, it provides limited taxonomic and functional resolution due to its focus on a single marker gene. In contrast, metagenomics offers a more comprehensive approach by sequencing the entire genetic material of a sample without targeted amplification^7^. This leads to a higher taxonomic resolution, enabling the identification of species- and strain-level variations within microbial populations^8^. Indeed, current metagenomic approaches have revealed more specific microbial biomarkers of health and disease across diverse populations and disease states, particularly in the context of inflammatory bowel disease and colorectal cancer ^9–12^. One recent notable example was the discovery of a distinct subclade of the species *Fusobacterium nucleatum* found to be enriched in colorectal cancer patients^13^. Furthermore, by characterizing the encoded genes and predicting their associated metabolic pathways, metagenomics can also uncover the functional potential of the microbiome community. For instance, prior metagenomic studies revealed that microbial pathways such as lipopolysaccharide biosynthesis and iron transport robustly distinguish diseased individuals from healthy controls across multiple conditions^14^.

However, developments in metagenome assembly and binning have revealed the existence of thousands of uncultured species in the human gut microbiome that still remain poorly understood^15–17^. These efforts culminated in the generation of the Unified Human Gastrointestinal Genome (UHGG) catalog^18^, a sequence database compiling over 3,000 uncultured bacterial species representing a large diversity of taxa that cannot yet be experimentally characterized. This now offers unprecedented opportunities for accurate, high-resolution metagenomic analysis to perform a dedicated investigation of the role of the uncultured microbiome in health and identify priority candidates for developing targeted culturing approaches.

Here we leveraged a dataset of over 11,000 human gut metagenomic samples across 13 noncommunicable diseases and healthy individuals to explore the clinical relevance of the uncultured gut microbiome. We discovered that the largely uncultured CAG-170 genus represents one of the strongest biomarkers of health and includes several putative keystone species distributed across healthy populations worldwide. Our study shows how genome-resolved metagenomics can shed light into the significance of novel, uncultured members of the gut microbiome in host health.

## Results

### Uncultured bacterial species are widespread across health and disease

To explore the clinical relevance of the uncultured microbiome in human health, we first gathered 8,672 human gut metagenomic samples across 36 studies, distributed among 24 countries and 13 noncommunicable diseases (Fig. 1 and Supplementary Table 1). These included 4,358 samples (50.3%) from healthy controls, 1,682 samples (19.4%) from patients with gastrointestinal disorders (adenoma, colorectal cancer, Crohn’s disease and ulcerative colitis), 1,215 (14%) from metabolic diseases (obesity, type 1 and type 2 diabetes), 1,099 (12.7%) from neurological disorders (myalgic encephalomyelitis, multiple sclerosis and Parkinson’s disease), and 318 samples (3.7%) from other diseases such as ankylosing spondylitis, bone disease, and rheumatoid arthritis. In terms of geographical breadth, samples were primarily collected from Europe (44%, *n* = 3,845), North America (37%, *n* = 3,230) and Asia (32%, *n* = 2,755), with most samples coming from the United States of America (37%), China (15%) and Denmark (8%). After quality-filtering and excluding samples with <500,000 reads, metagenomic samples represented a median of 25.8 million reads (interquartile range, IQR = 9–46 million reads, Extended Data Fig. 1a).

**Figure 1.**
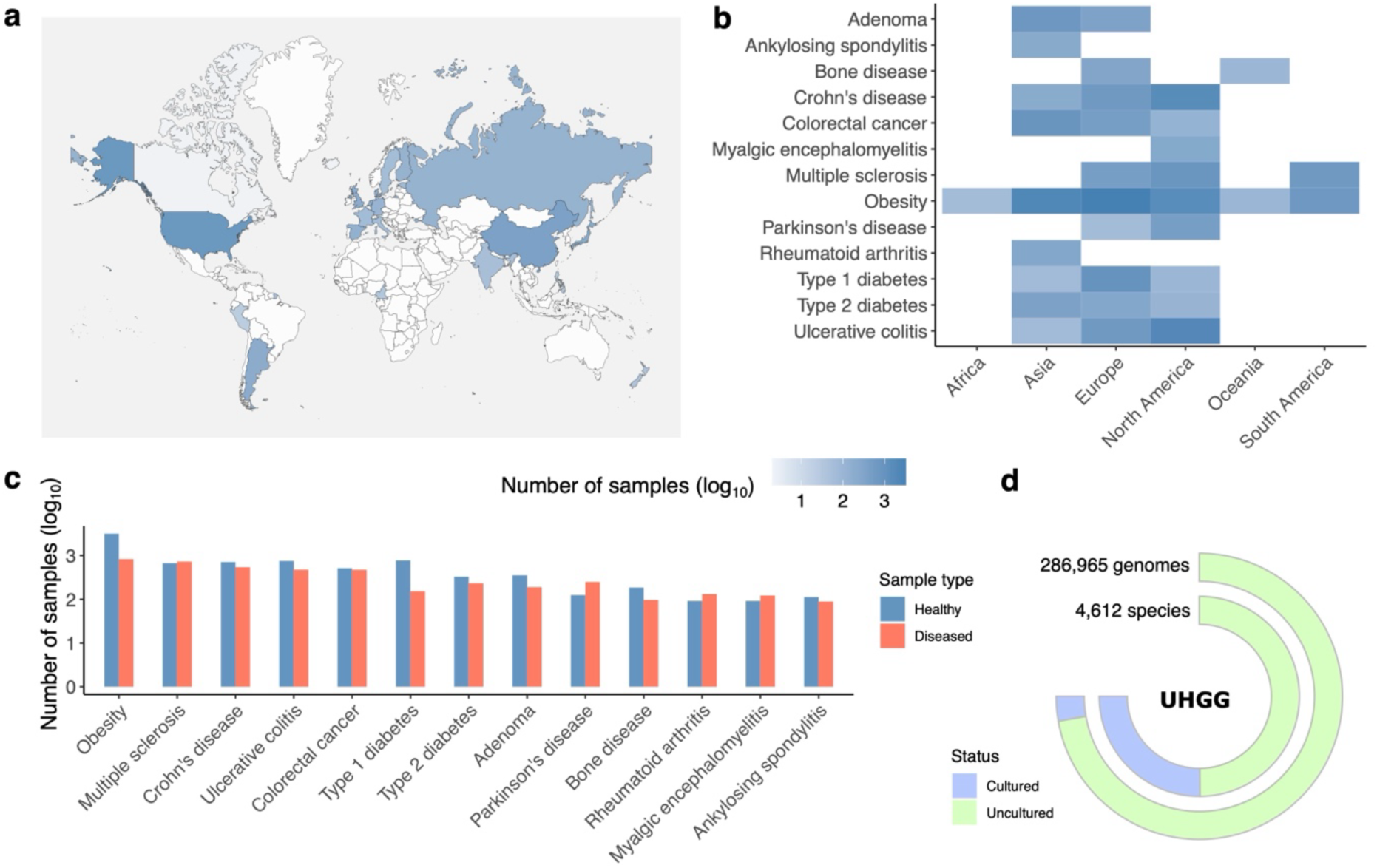
Meta-analysis of the uncultured microbiome in health and disease. **a**, Geographic distribution of the case-control metagenomic samples analysed in this study. **b**, Heatmap denoting the number of samples (log_10_-transformed) processed per disease and per continent. **c**, Number of healthy and case (disease) metagenomic samples analysed per condition. **d**, Distribution of the number of cultured (blue) and uncultured (green) genomes and species present in the Unified Human Gastrointestinal Genome (UHGG) catalog.

Metagenomic samples were individually mapped to the Unified Human Gastrointestinal Genome (UHGG) catalog^18^ — a comprehensive reference database of genomic sequences from currently known prokaryotic species within the human gut microbiome. The UHGG includes 4,612 species, of which 66.5% (*n* = 3,067) belong to prokaryotic species currently without a representative isolate genome (hereafter referred to as ‘uncultured’ species; Fig. 1d). To accurately quantify the prevalence and abundance of each species in each sample, we employed a genome-based mapping approach that integrates metrics of both observed and expected genome breadth, along with depth of coverage, using previously validated thresholds^19^. Compared to methods based solely on marker genes, genome-based approaches that apply minimum genome breadth filters offer improved accuracy for species detection^8^. We obtained a median read mapping rate of 85.6% (IQR = 83.5–87.4%) across all samples, which reduced to 61% (IQR = 57.4–64.2%) after applying further filters based on breadth and depth of coverage (Extended Data Fig. 1b; see Methods for further details).

A median of 187 prokaryotic species was detected per sample (IQR = 92–313 species), with 30.7% (IQR = 22–42%) belonging to uncultured bacteria. A total of 1,209 uncultured species were detected in at least 1% of samples, with the most prevalent species belonging to the genera *Faecalibacterium*, CAG-103, and CAG-170 (Extended Data Fig. 2a), the latter two representing genera currently without a cultured type strain. The proportion of uncultured species was generally consistent across health states (Extended Data Fig. 2b). However, in the context of Crohn’s disease, ulcerative colitis and bone disease, uncultured species were significantly more prevalent in healthy controls compared to disease samples (Wilcoxon rank-sum test, False Discovery Rate, FDR < 0.05). Regarding overall abundance, uncultured species represented a median of 11.8% of mapped reads (Extended Data Fig. 2b), indicating that although uncultured bacteria comprise approximately 30% of the species in the human gut, their relative abundance is generally lower compared to cultured genomes.

Beyond calculating the number of species per sample, we estimated alpha diversity metrics using the Shannon index to account for both species richness and evenness. Based on the abundance patterns of the uncultured species detected, healthy controls showed significantly higher alpha diversity (linear mixed-effects model, FDR < 0.05) when compared to subjects from four diseases (ankylosing spondylitis, Crohn’s disease, obesity, and ulcerative colitis; Extended Data Fig. 3a). Although the alpha diversity of cultured and uncultured bacteria generally correlated (Pearson R² = 0.45, *P* < 0.0001; Extended Data Fig. 3b), we detected statistically significant differences between the diversity of ulcerative colitis patients and healthy controls only when using the uncultured abundance profiles. Overall, these results show that uncultured bacteria are widely distributed across diverse health states; are disproportionately prevalent among certain healthy populations; and their diversity can better discriminate case and control samples in the context of ulcerative colitis.

### The uncultured microbiome is clinically relevant

Given the potential of the gut microbiome for disease prediction and diagnostic applications^20–22^, we compared how well the composition of the cultured and uncultured microbiome could differentiate between healthy and diseased individuals. We evaluated three supervised machine learning approaches — ridge regression, random forest, and gradient boosting — across the 13 noncommunicable diseases tested, and found that the choice of method was significantly associated with model performance (Kruskal-Wallis test, *P* = 0.009; Extended Data Fig. 4a). When using the uncultured microbiome alone, ridge regression yielded the highest Area Under the Receiver Operating Characteristic Curve (AUROC) in 8 out of 13 diseases and a median AUROC of 0.728 (Extended Data Fig. 4b). This was followed by gradient boosting, which had the highest AUROC in 3 diseases, and random forest, which outperformed in 2 diseases. Consequently, all subsequent analyses were conducted using ridge regression to facilitate further comparisons.

On a per-disease basis (pooling subjects across all studies of one disease), models based on the uncultured microbiome performed best for ankylosing spondylitis (median AUROC = 0.916), ulcerative colitis (median AUROC = 0.814), and Crohn’s disease (median AUROC = 0.778; Extended Data Fig. 5a). Overall, the performance of models using uncultured versus cultured species was generally comparable, with a median AUROC difference (AUROC_uncultured_ – AUROC_cultured_) of −0.01, and no statistically significant differences observed for most diseases (7 out of 13 diseases; Extended Data Fig. 5a). Interestingly, for adenomas, models with uncultured species performed significantly better (Wilcoxon rank-sum test, FDR = 0.034) compared to those with cultured species. Therefore, despite the uncultured microbiome representing only ∼12% of the microbial abundance detected in each sample, it offers comparable performance for disease classification compared to models based on cultured species alone.

We further investigated the generalizability of the machine learning models trained on both cultured and uncultured species using a cross-validation approach across different studies of the same disease and between different diseases (Extended Data Fig. 5b). When analysing different studies from the same disease, gut-associated disorders (Crohn’s disease, ulcerative colitis and colorectal cancer) showed the greatest generalizability (≥0.7 AUROC in the cross-study validation), with type 1 diabetes, bone disease and Parkinson’s disease exhibiting the lowest performance (<0.5 AUROC). Furthermore, in a cross-disease classification analysis, machine learning models were generally found to be disease specific, with limited cross-disease transferability (Extended Data Fig. 5c). However, a few notable exceptions were observed. Moderate classification performance (AUROC ≥0.7) was observed when models trained on myalgic encephalomyelitis were applied to Crohn’s disease samples; when colorectal cancer models were tested on Crohn’s disease; and when ulcerative colitis models were used to classify ankylosing spondylitis samples or vice-versa. Interestingly, Crohn’s disease is considered a risk factor for both myalgic encephalomyelitis^23^ and colorectal cancer^24^, while ulcerative colitis and ankylosing spondylitis are inflammatory disorders that share genetic predispositions and overlapping pathophysiological mechanisms^25^. These findings suggest that gut-associated disorders such as Crohn’s disease, ulcerative colitis and colorectal cancer have shared microbial features across different populations and diseases that may represent more general markers of health and disease.

### The genus CAG-170 is the strongest uncultured biomarker of health

To identify robust microbial biomarkers of health and disease we performed a differential abundance analysis using two complementary statistical methods based on generalized linear models^26,27^ (see Methods). After accounting for the subject’s age group, continent of origin, sample read depth and batch (that is, study source), we identified 715 prokaryotic species associated with at least one disease (398 cultured, 317 uncultured; Fig. 2a and Supplementary Table 2). The number of biomarkers varied by disease, with most species found to be associated with Crohn’s disease (*n* = 414), ulcerative colitis (*n* = 396), obesity (*n* = 227) and/or colorectal cancer (*n* = 60). Given the effect size and direction of association across the 13 diseases tested, we calculated a combined effect size to denote the overall strength of association of each species with either health or disease. A total of 342 species were classified as disease biomarkers (higher abundance in disease; Extended Data Fig. 6a), whereas 373 species were health biomarkers (higher abundance in health; Extended Data Fig. 6b). Of note, 74% of the disease biomarkers were uniquely associated with one disease, whereas 56% of the health biomarkers were consistent across multiple diseases, suggesting a more conserved signal of health. Only 25 species (3%) displayed conflicting associations — linked to health in some disease cohorts and to disease in others — indicating that most species exhibit consistent associations across different conditions.

**Figure 2.**
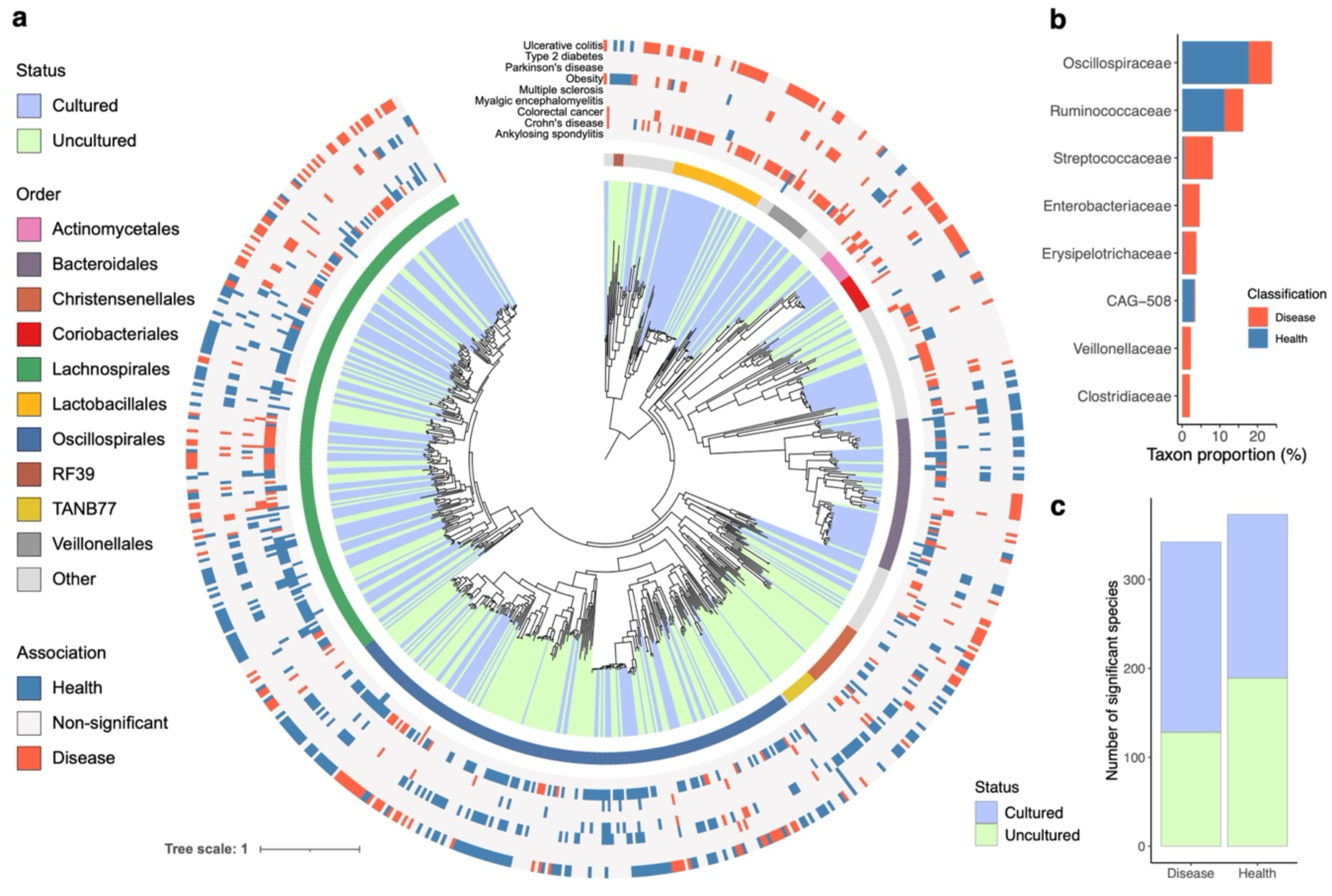
Phylogenetic structure of health and disease biomarkers. **a**, Phylogenetic tree of the 713 bacterial biomarkers identified as associated with either health or disease. Outermost layers denote the association of each species across different conditions. Innermost layer indicates the taxonomic affiliation (order rank). Clades are coloured based on the cultured status of the species. **b**, Bacterial families found to be significantly associated with either health or disease (Fisher’s Exact Test, FDR < 0.05). Taxon proportion represents the fraction of species among the predicted biomarkers that were linked to either health or disease within each family. **c**, Number of cultured and uncultured species found as either health- or disease-associated. Proportion of uncultured species was found to be significantly higher (χ^2^-test, *P* = 0.00049) among the health biomarkers.

At a taxonomic level, the families Oscillospiraceae, Ruminococcaceae and CAG-508 were found to be significantly overrepresented among the health biomarkers (Fisher’s Exact Test, FDR < 0.0001; Fig. 2b), whereas Streptococcaceae, Enterobacteriaceae, Erysipelotrichaceae, Veillonellaceae and Clostridiaceae were overrepresented in disease. This signal was found to be consistent at higher taxonomic ranks, with the order Oscillospirales most strongly associated with health, and the order Lactobacillales most overrepresented in disease.

We discovered that the proportion of uncultured species was significantly higher (χ^2^-test, *P* = 0.00049) among the health-associated biomarkers compared to the disease-associated species, implicating the uncultured microbiome as a potential marker of health (Fig. 2c). To investigate this further we aimed to identify the specific uncultured taxa that most strongly underlined a healthy state. We therefore ranked all bacterial genera present in the UHGG based on a combined measure that incorporated three key metrics: (i) the total number of health-associated species (biomarkers) detected per genus; (ii) a genus-level effect size normalized across diseases and species; and (iii) the proportion of uncultured species currently known within each genus (Fig. 3a and Supplementary Table 3). Using this combined measure, we determined that the unclassified genus CAG-170 was the top-ranking taxon (Fig. 3b). This result was found to be consistent when aggregating effect sizes using an independent meta-analysis approach (Extended Data Fig. 7; see Methods for further details). In fact, CAG-170 was the only genus that matched all the following criteria: (i) >10 significant health biomarkers; (ii) absolute effect size >0.5; and (iii) >75% uncultured species (Supplementary Table 3). The taxon CAG-170 is a largely unknown bacterial genus from the Oscillospiraceae family, represented by 1,046 genomes and 13 species in the UHGG. Within our dataset, CAG-170 species were detected at a median prevalence of 20.3% (IQR = 13.4–29.1%) and a median relative abundance of 0.07% (IQR = 0.03-0.18%; Fig. 3c). In the differential abundance analysis, 11/13 (85%) of CAG-170 species were found to be associated with health in the context of Crohn’s disease, myalgic encephalomyelitis, ulcerative colitis and obesity (Fig. 3c). Collectively, these results show that CAG-170 has a consistent health-associated signal across multiple conditions and multiple species.

**Figure 3.**
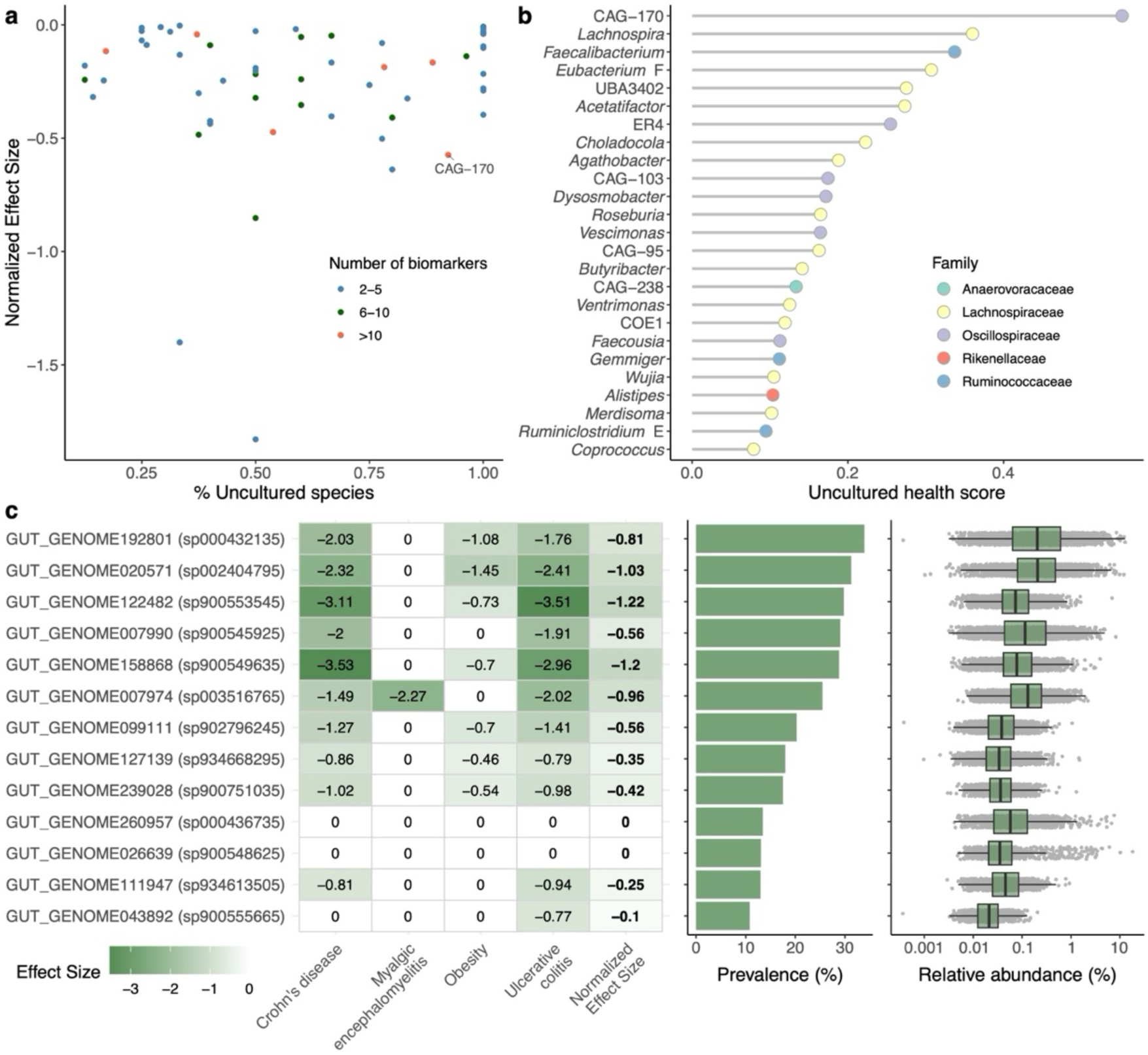
CAG-170 is the most significant uncultured genus in health. **a,** Proportion of uncultured species relative to the estimated effect size of each bacterial genus in the UHGG. A negative effect size indicates an association with health. Each genus is coloured according to the number of health biomarkers identified (only genera with an overall association with health are shown). **b**, Bacterial genera ranked by the absolute value of their uncultured health score, calculated as the product of the number of significant species (log_10_-transformed), normalized effect size, and proportion of uncultured species. **c**, Effect sizes, prevalence, and relative abundance of the 13 species of CAG-170 present in the UHGG. In the boxplots, the centre line represents the median. Whiskers extend to the furthest point within 1.5 times the IQR from the box. The number of data points in the boxplots corresponds to the species prevalence among 8,672 samples.

### CAG-170 is a candidate keystone genus of the healthy microbiome

Having identified uncultured health-associated biomarkers through the analysis of case-control cohorts, we sought to define the broader characteristics of the healthy uncultured microbiome on a global scale. To achieve this, we aggregated all healthy control samples from the case-control studies analysed in this work (*n* = 3,614) and incorporated an independent dataset comprising an additional 2,443 metagenomes from healthy individuals (Supplementary Table 1). We first examined prevalence patterns on a per-continent basis (across Africa, Asia, Europe, North America, Oceania and South America) to assess the presence of a dominant core microbiome either within or across regions. Cultured species were found to be the most prevalent in Asia (median prevalence = 9.7%; IQR = 3.3–31%), whereas uncultured species had their highest prevalence in Africa (median = 9.2%; IQR = 1.7–23.1%). Across the full spectrum of gut species, we identified 29 cultured and 2 uncultured species within the top 1% most prevalent species (>70% prevalence) across the six continents analysed (here defined as the ‘core’ set of microbiome species; Fig. 4a). When averaging species prevalences across continents, only one cultured species (*Blautia wexlerae*) reached a mean worldwide prevalence >90% among healthy individuals (Fig. 4a), further emphasizing that most microbiome species are not universally conserved across diverse global populations. The two core uncultured species belonged to the *Faecalibacterium* genus, and included an unclassified species detected in 77% of samples.

**Figure 4.**
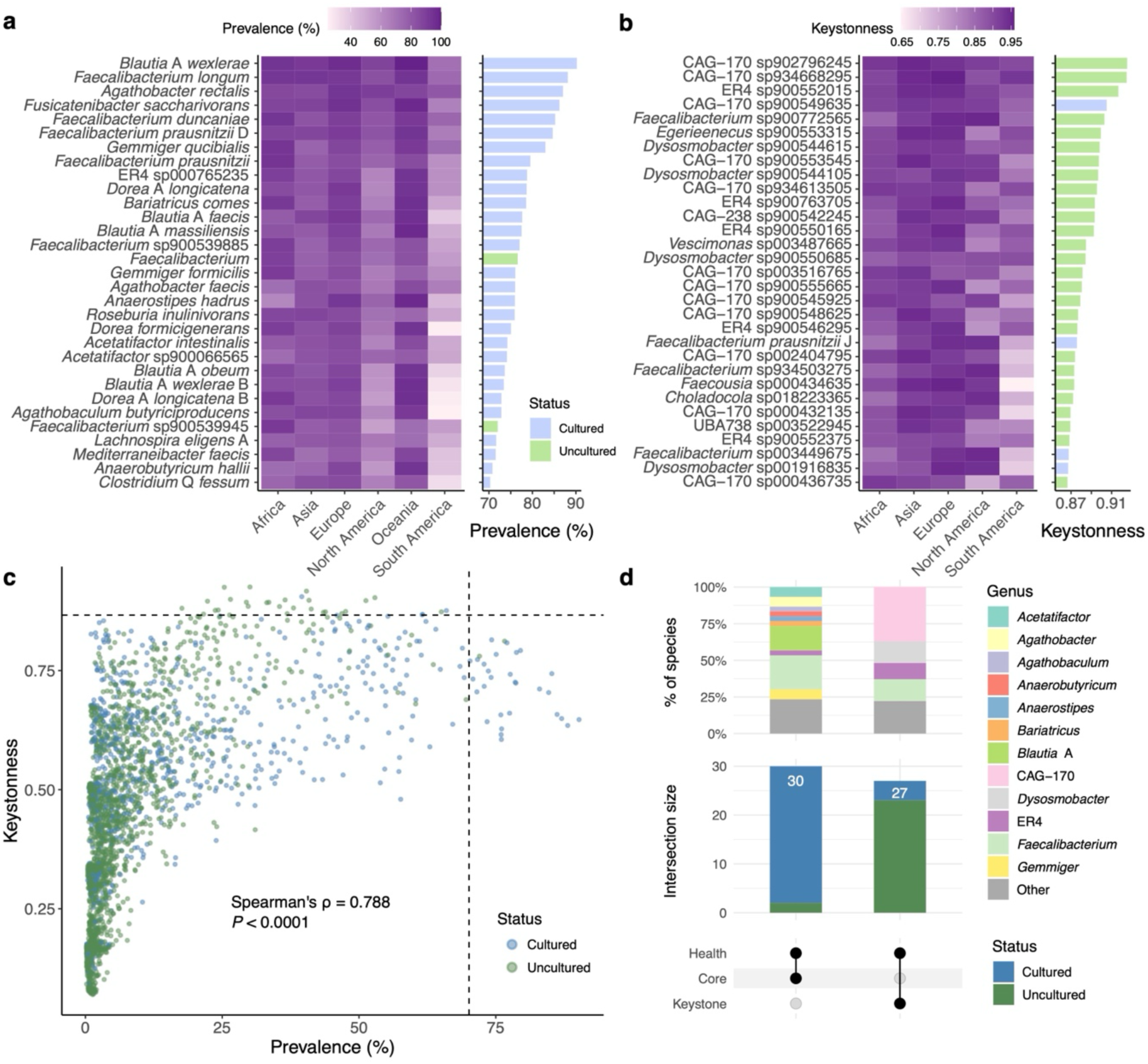
CAG-170 is the top candidate keystone genus among healthy microbiomes worldwide. **a,** Prevalence of the top 1% most common species (defined as ‘core’). The heatmap shows prevalence across continents, and the bar plot summarizes mean prevalence. **b**, Keystonness scores of the top 1% most central species (defined as ‘keystone’). The heatmap shows continent-specific values, and the bar plot shows the mean keystonness across continents. **c**, Distribution of mean prevalence and keystonness scores for cultured and uncultured species. Dashed lines mark the thresholds we defined for core and keystone species. **d**, Overlap among species classified as health biomarkers, core, and keystone (only species that overlapped in at least two groups are shown). Vertical bars indicate the number of species in each intersection, with stacked bars above showing their taxonomic affiliation (genus level).

To complement the prevalence analysis, we further explored putative ecological interactions of the healthy microbiome by modelling interspecies co-abundance patterns. Similar to the prevalence analysis, we constructed co-abundance networks separately for samples from each continent. From these networks, we computed three centrality measures for each species — closeness, betweenness, and degree — to capture different aspects of their influence on the overall network structure. It has been previously shown that a combination of high degree, high closeness centrality and low betweenness centrality derived from co-occurrence networks can identify keystone taxa with 85% accuracy^28,29^. Accordingly, we integrated these metrics into a composite keystonness score averaged across continents, and defined as candidate keystone species those ranking within the top 1% of this combined measure. This revealed a total of 31 keystone species, of which 27 were classified as uncultured (Fig. 4b). Importantly, the CAG-170 genus was the most represented taxon among this set, with 12 species classified as putative keystone. Even after filtering network edges with two different correlation thresholds, CAG-170 remained the most frequent keystone genus identified (Extended Data Fig. 8).

Lastly, combining all these results we assessed the overlap among species classified as health-associated, disease-associated, core, and keystone to determine whether there were consistent patterns (Supplementary Table 4). Although prevalence and keystonness was found to generally correlate (Spearman’s ⍴ = 0.788, *P* < 0.0001; Fig. 4c), there was no overlap between species that were classified as core and keystone (Fig. 4d). This suggests that the most prevalent microbial species are not necessarily the ones with the strongest ecological impact. With regards to health state, most core and keystone species were identified as health biomarkers (Fig. 4d), suggesting that such species are typically depleted in disease states. Uncultured bacteria were more frequent among species that were both keystone and health-associated, with the genus CAG-170 again observed as the most represented within this group (Fig. 4d). Together, these findings highlight that the uncultured microbiome — particularly species from the CAG-170 genus — may play key ecological roles within the global healthy microbiome.

### Species from CAG-170 have a unique functional capacity

To go beyond a taxonomic characterization of the microbiome across health states, we examined which functions were enriched among health- or disease-associated species. By profiling the KEGG Orthologs^30^ (KOs) encoded in each health or disease biomarker, we identified 46 KOs specifically enriched among the health-associated species, and 146 KOs among the disease-associated biomarkers (logistic regression, FDR < 0.0001; Supplementary Table 5). Among these candidate KOs, we observed that functions related with cell motility and immune regulation (e.g., flagellar biosynthesis proteins) were most strongly associated with health; whereas those involved in osmotic and oxidative stress were enriched among the disease biomarkers (Extended Data Fig. 9). Flagellar proteins from commensal gut bacteria, particularly flagellin, have been shown to support health through immune regulation — such as activation of Toll-like receptor 5 — which in turn can help strengthen gut barrier integrity and prevent pathogen colonization^31^. In contrast, the presence of oxidative and osmotic stress resistance genes among disease-associated species may facilitate their survival under inflammatory and stressful conditions of the diseased gut environment^32^.

Given the strong health-associated signal we identified for species within the CAG-170 genus, we additionally performed a dedicated analysis of its functional repertoire to pinpoint features that distinguish CAG-170 from closely related bacterial clades. To this end, we analysed all high-quality (>90% completeness) and non-redundant genomes from the Oscillospiraceae family in the UHGG collection (*n* = 4,025 genomes from 152 species; Fig. 5a). This dataset spanned 28 genera and was predominantly represented by genomes belonging to ER4 (*n* = 1,210), *Faecousia* (*n* = 522) and *Vescimonas* (*n* = 445), with 332 genomes assigned specifically to the CAG-170 genus. We applied a mixed-effects logistic regression model to compare the presence of KEGG Orthologs (KOs) in CAG-170 genomes versus all other Oscillospiraceae genera. This analysis identified 135 KOs significantly enriched and 141 KOs significantly depleted in CAG-170 genomes (FDR < 0.05; Fig. 5b and Supplementary Table 6). Enriched genes spanned diverse metabolic pathways, most notably including 14 KOs covering the cobalamin (vitamin B12) biosynthesis pathway (Fig. 5c). They also encompassed genes involved in methionine and serine biosynthesis (*metA, metY, serA*), stress tolerance and DNA repair functions (*mutL, mutY, uvrD/pcrA, dinG*), sporulation-related genes (*spoVFA, spoVFB, spoVG*), and broad nutrient uptake systems (e.g., multiple sugar transporters *msmK/msmX, rbsB*). In contrast, depleted genes were linked to pro-inflammatory or virulence-associated functions, including lipopolysaccharide/rhamnose biosynthesis (*rfbB, rfbC, rfbD, rmlB, rmlC, rmlD*), spore germination proteins (*cwlJ, sleB*), and metal stress regulators (*cadC/smtB*). Several vitamin and nucleotide biosynthesis genes were also missing, such as those in the thiamine (*thiD, thiE, thiM, tenA)* and pantothenate (*panE*) pathways. Notably, some of the most strongly depleted genes among CAG-170 genomes are involved in key steps of the arginine biosynthesis pathway (*carA, carB, argC, argJ*; Fig. 5c), indicating likely arginine auxotrophy and dependence on host or microbial cross-feeding. Coincidentally, the only reported successful cultivation of one CAG-170 isolate involved arginine supplementation in the culture medium^33^ — an approach not employed in earlier culturing attempts^3,34^ — suggesting it may be important for the *in vitro* growth of CAG-170. Collectively, these findings show that an enrichment of vitamin B12 biosynthesis and depletion of arginine production are key traits of CAG-170 species. These new insights will open opportunities to obtain a more comprehensive mechanistic understanding of this understudied genus.

**Figure 5.**
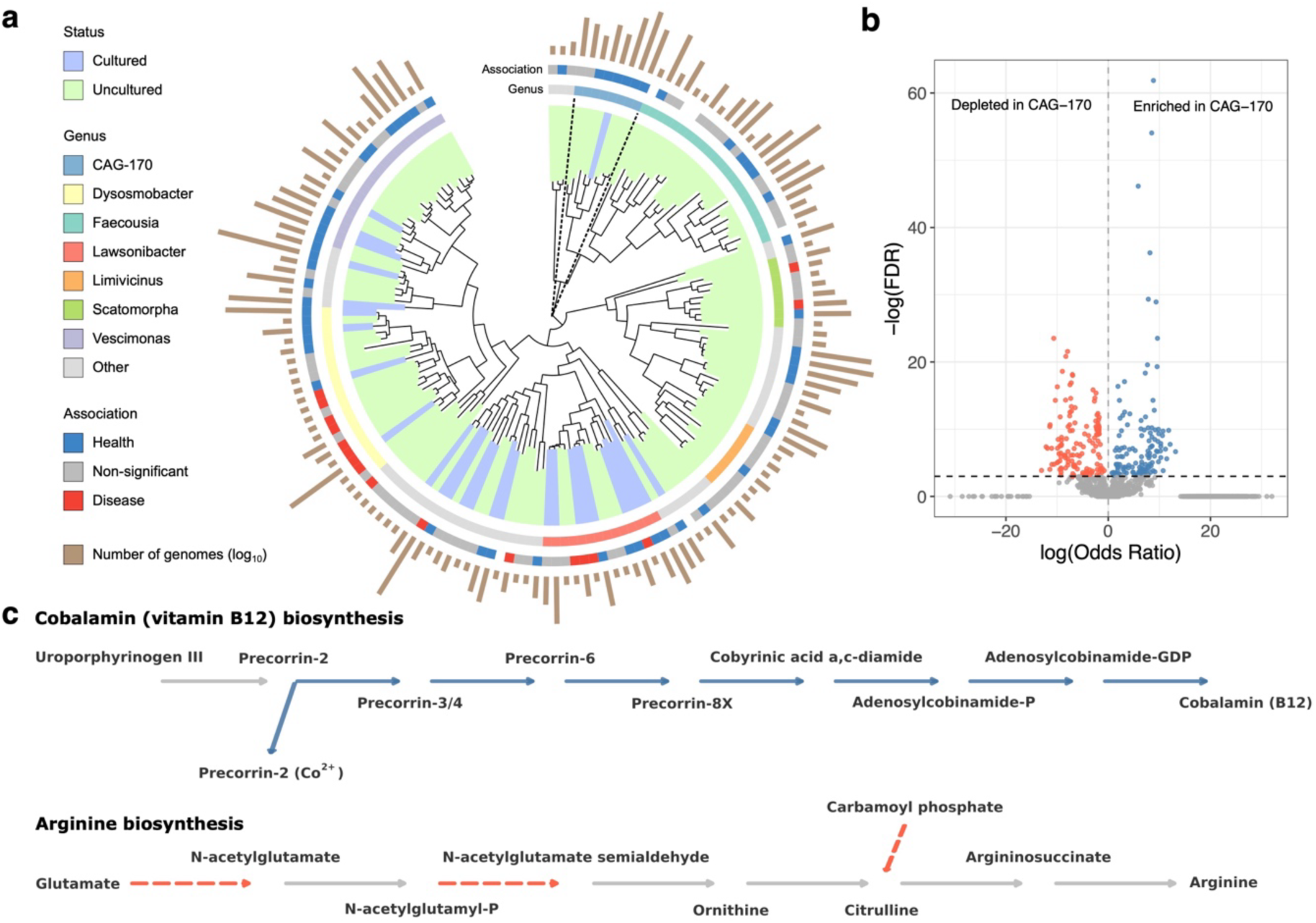
Phylogenetic and functional analysis of CAG-170. **a,** Phylogenetic tree of 152 species from the Oscillospiraceae family, with clades and outer layers labelled by species culturing status, genus affiliation and association with health state. Bar plot in the outermost layer denotes the log_10_-transformed number of high-quality (>90% complete) genomes of the corresponding species present in the UHGG catalog. The clade belonging to the CAG-170 genus is highlighted with a dashed outline. **b**, Volcano plot showing the KEGG Orthologs (KOs) significantly enriched (positive effect size) or depleted (negative effect size) among CAG-170 genomes compared to other Oscillospiraceae genera. Adjusted *P* values (FDR) were derived from a mixed-effects logistic regression. **c,** Simplified diagram of the biosynthetic pathways for cobalamin (vitamin B12) and arginine production, focusing on reaction steps detected as significantly enriched (solid blue) or depleted (dashed red) in CAG-170.

### CAG-170 abundance and subspecies diversity in a longitudinal cohort

Given that cross-sectional case-control cohorts cannot capture microbiome dynamics over time, we further characterized the abundance and subspecies diversity of CAG-170 within a longitudinal cohort of inflammatory bowel disease (IBD) patients and controls^9^. This cohort was recruited as part of the second phase of the Human Microbiome Project (HMP2), characterizing a total of 132 Crohn’s disease, ulcerative colitis and non-IBD individuals across a year, sampled every 2 weeks. Each microbiome sample had an associated ‘dysbiosis’ score, measuring the extent of deviation from a healthy gut microbiome composition inferred in the original study^9^. By reanalysing this dataset with the UHGG catalog, we found that the abundance of CAG-170 negatively correlated with an individual’s dysbiosis score across time (linear mixed-effects model, *P* < 0.0001; Fig. 6a). This result was also consistent after classifying samples into either low or high dysbiosis, showing that the abundance of CAG-170 was significantly higher in low dysbiosis samples (linear mixed-effects model, *P* = 6.11 ⨉ 10^-12^; Fig. 6b). This supports the notion that there is a proportional negative relationship between the abundance of CAG-170 and the extent of microbial imbalance in the gut microbiome.

**Figure 6.**
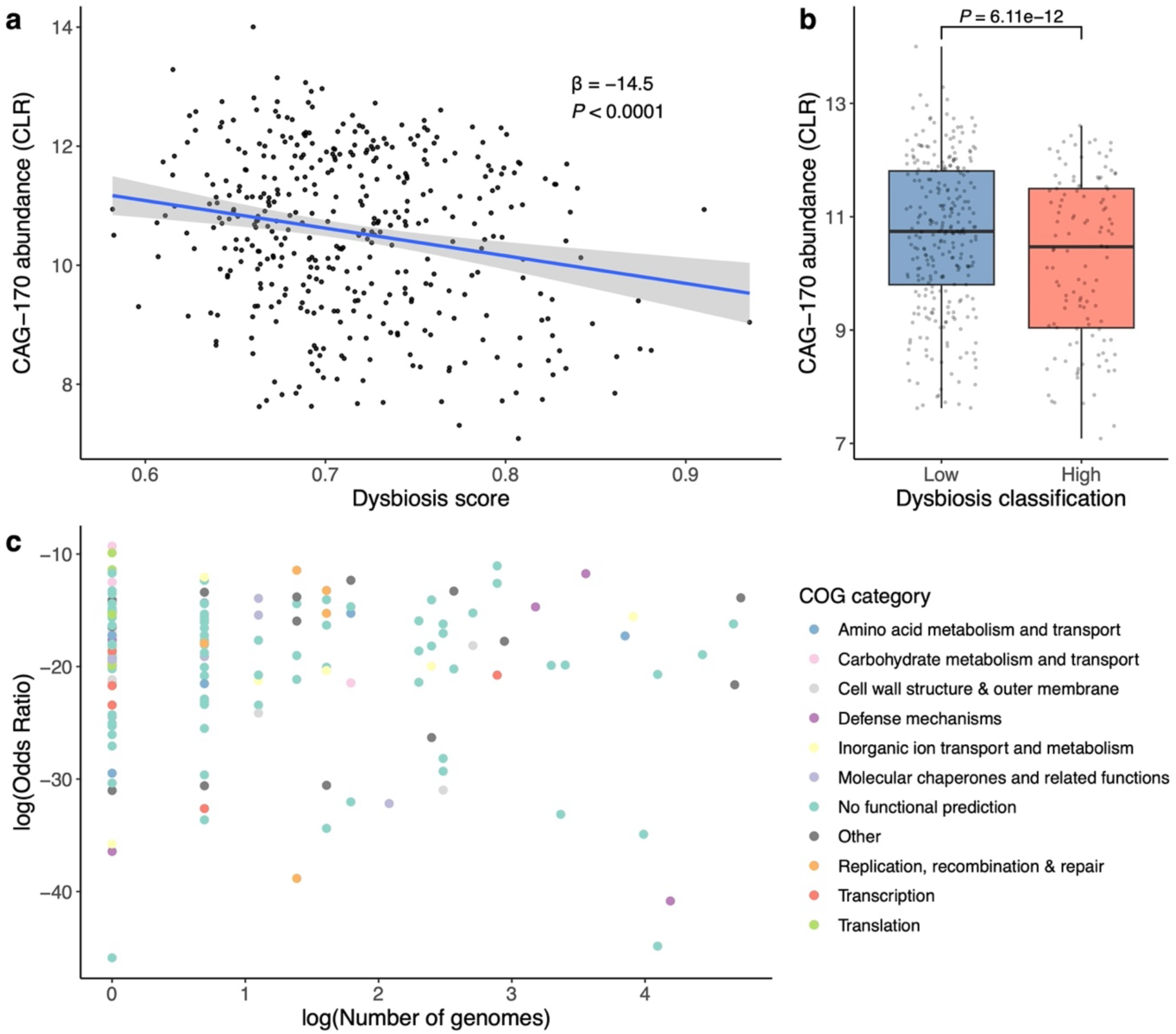
The abundance and subspecies diversity of CAG-170 correlates with health. **a,** Correlation between the abundance of CAG-170 (converted to centred log-ratio) with the dysbiosis score estimated for each sample from the HMP2-IBD study. **b**, Comparison of CAG-170 abundance when stratifying samples (*n* = 1,118) into ‘low’ or ‘high’ dysbiosis based on the median value. For visualization purposes only, samples with no CAG-170 species were excluded. Within each boxplot, the centre line represents the median. Whiskers extend to the furthest point within 1.5 times the IQR from the box. **c**, Representation of the 160 CAG-170 genes negatively associated with dysbiosis, based on the number of high-quality genomes containing each gene and the corresponding effect size (log odds ratio). Genes are coloured by their COG functional category. Effect sizes and *P* values were calculated using a mixed-effects linear model that accounted for repeated measures within subjects.

Next, to move beyond abundance- and species-level analyses, we assessed whether the subspecies genetic diversity of CAG-170 was associated with health and lower dysbiosis scores. We first generated pan-genomes of each of the 13 CAG-170 species, identifying a total of 3,589 unique accessory genes (1,186 of which detected in >1% of the samples in the HMP2 cohort). By investigating the gene prevalence patterns among the HMP2-IBD samples, we found 160 CAG-170 accessory genes negatively associated with dysbiosis score (FDR < 0.05; Fig. 6c and Supplementary Table 7), and no genes with a positive association. These health-associated genes covered various functional categories, including transport and nutrient acquisition (e.g., ABC transporters, ion pumps and efflux systems); stress response and cellular maintenance (e.g., chaperones, DNA repair and sporulation); and information processing (e.g., translation factors and transcriptional regulators). However, 87 candidate genes (54%) had no recognized function, representing a large uncharacterized diversity within CAG-170 that remains unexplored. Overall, these results highlight CAG-170 as a key uncultured taxon in gut health, with both its abundance and genetic diversity consistently linked to a low dysbiosis, health-associated microbiome.

## Discussion

Here we provide a comprehensive meta-analysis of the human gut uncultured microbiome across a broad spectrum of diseases, populations, and geographic regions. By integrating large-scale metagenomic data with genome-based species profiling, machine learning, ecological modelling, and functional prediction, we reveal that uncultured bacterial species are widespread, clinically relevant, and carry unique ecological and functional signatures.

Despite representing only ∼30% of species and ∼12% of the total microbial abundance per sample, uncultured bacteria were found to be strong biomarkers of microbial diversity and health across multiple datasets. We demonstrated that the performance of classifiers based on uncultured species generally matched — and in one case outperformed — models trained on cultured species alone. This highlights the diagnostic potential of these underrepresented taxa, even when present at low abundance. Among the uncultured bacteria identified, health-associated species were more commonly conserved across multiple diseases, whereas disease-associated species tended to be disease specific. These findings suggest that the Anna Karenina principle may also extend to the microbiome: healthy microbiomes are more similar to one another, whereas diseased microbiomes are more individualized and variable. Our study expands on previous efforts investigating the unique properties of the healthy microbiome^35,36^ by providing new insights specifically from uncultured gut bacteria.

Our most notable finding was the identification of the uncharacterized genus CAG-170 as a consistent and robust biomarker of gut health. This was supported by three key results: 1) CAG-170 was the uncultured genus most significantly enriched in healthy individuals; 2) CAG-170 was more frequently represented among candidate keystone species in healthy populations than any other genus — cultured or uncultured; and 3) its abundance and genetic diversity correlated with microbiome dysbiosis over time in a longitudinal IBD cohort. Taken together, this suggests that CAG-170 may play a key ecological role in shaping and maintaining a stable gut microbial community in healthy individuals.

Functional characterization of CAG-170 genomes revealed unique biological attributes, including an enrichment of various genes related to vitamin B12 biosynthesis, when compared to other species from the Oscillospiraceae family. In the gut microbiome, vitamin B12 serves as a cofactor for microbial enzymes leading to short-chain fatty acid production and amino acid metabolism^37–39^. Therefore, a higher capacity for vitamin B12 biosynthesis may partly underly the health-associated role of CAG-170. Furthermore, we also detected a depletion of pro-inflammatory genes and those involved in arginine biosynthesis among CAG-170 species, the latter of which could be related with the difficulty in culturing this genus. As a result, CAG-170 remains largely understudied. In fact, since nearly all available CAG-170 genomes are metagenome-assembled, 16S rRNA gene sequences for this group are also underrepresented in established databases such as SILVA^40^. This reflects a broader challenge in microbiome research: potentially relevant and ecologically important taxa remain invisible to traditional culturing techniques and certain marker gene surveys.

While we analysed a large and diverse range of diseases and populations, it is important to note some of the limitations of our study. First, the analysis of several diseases (especially of understudied conditions) may have been underpowered to detect subtle microbiome associations, particularly when analysing low-abundance, uncultured species. Secondly, our results are based on statistical associations and do not establish direct microbiome-disease causality, which requires longitudinal interventional studies and mechanistic validation using *in vivo* models. However, the lack of available isolates for these uncultured species currently limits further experimental work. Expanding cultivation efforts with targeted media supplementation, alongside continued development of genome-resolved metagenomics, will be essential for closing these knowledge gaps.

In conclusion, our findings provide important insights into the uncultured gut microbiome as a fundamental and underappreciated component of human health. As microbiome research continues to evolve, future efforts should prioritize the integration of metagenomics, culturing, and experimental validation to characterize these hidden, yet ecologically significant microbial species. This will not only improve our understanding of the role of the microbiome in health and disease but may also inform the development of next-generation microbial therapeutics that leverage the full diversity of the gut ecosystem.

## Acknowledgements

The authors thank all members of the Microbiome Function and Diversity group for helpful feedback and suggestions. Funding was provided by a Career Development Award from the Medical Research Council (MR/W016184/1) to A.A. This project was also supported with funding from the Cambridge Centre for Data-Driven Discovery and Accelerate Programme for Scientific Discovery, made possible by a donation from Schmidt Sciences.

## Author contributions

A.C.S. curated the sample metadata; performed metagenomic analyses and helped write the manuscript draft. J.L. performed the subspecies genetic analysis of CAG-170. Q.Y. assisted in the metagenomic analyses and data curation. E.M. co-supervised the work and contributed to the subspecies and statistical analyses of CAG-170. A.A. supervised the work, performed metagenomic analyses and wrote the manuscript draft. All authors read, edited and approved the manuscript.

## Competing interests

The authors declare no competing interests.

## Methods

### Metagenomic data collection and preprocessing

In this study, we collected 11,115 human gut metagenomic samples from the European Nucleotide Archive (ENA)^41^, spanning 62 studies and 39 countries. This dataset included 8,672 metagenomes from 36 case-control studies of 13 noncommunicable diseases, as well as an additional 2,443 samples from 26 studies composed exclusively of healthy individuals (Supplementary Table 1). Sample metadata were extracted and manually curated by reviewing the associated publications and supplemented with information from the curatedMetagenomicData^42^ and GMrepo^43^ databases. Samples were selected based on the following criteria: (1) availability of metadata on health status, age group, and country of origin; (2) absence of diagnosed acute infections; and (3) no reported antibiotic use within the previous month. Raw metagenomic datasets were downloaded from ENA using fastq-dl (v.2.0.4; https://github.com/rpetit3/fastq-dl), followed by quality control using TrimGalore (v.0.6.0)^44^. Human DNA contamination was removed by aligning reads to the human reference genome (GRCh38) using BWA-MEM (v.0.7.16a-r1181)^45^. Samples with fewer than 500,000 paired-end metagenomic reads after filtering were excluded from further analysis.

### Generating a species-level genome database

To generate a custom database of genomes for metagenomic read mapping, we gathered all species representatives genomes from the Unified Human Gastrointestinal Genome (UHGG) catalog (v.1)^18^ and filtered those matching all of the following criteria, as previously described^19^: (1) singletons (that is, the respective species includes only one genome); (2) <90% genome completeness based on CheckM (v.1.0.11)^46^; and (3) classified by GUNC (v.1.0.3)^47^ as chimaeric (‘clade_separation_score’ >0.45, ‘contamination_portion’ >0.05 and ‘reference_representation_score’ >0.5). This removed 32 species, resulting in a total of 4,612 species representatives. The final curated database was indexed using BWA (v.0.7.16a-r1181, ‘bwa index’) for subsequent read mapping.

### Species classification and cultured status

Species classification based on their cultured status was determined according to two main criteria: (1) the genome type of the species representative (isolate or MAG); and (2) similarity to isolate genomes present in the NCBI RefSeq database (release 219). For this purpose, each genome was first screened against the RefSeq database using Mash (v.2.3)^48^ to identify its most similar relative (lowest Mash distance). Subsequently, DNAdiff (v.1.3)^49^ was used to compute whole-genome average nucleotide identities (ANI) between each pair. A species was considered ‘cultured’ if it was represented by an isolate genome or had >95% ANI over a minimum 30% alignment fraction to its closest RefSeq isolate. This classification was further refined through manual curation informed by literature searches, for example, when newly published UHGG species isolates were not represented in RefSeq. In addition, to account for taxonomy updates since the initial release of the UHGG, species representatives were reclassified with GTDB-Tk (v.2.3.2) (database release 214)^50^ using the ‘classify_wf’ with default parameters.

### Profiling species prevalence and diversity

Each quality-filtered metagenomic sample was mapped and quantified against our custom UHGG catalog, as described previously^19^. Briefly, metagenomic reads were first aligned using BWA-MEM and processed with SAMtools (v1.9)^51^. For each species in the reference database, we calculated the following metrics per sample: breadth of coverage, depth of coverage, total read counts and counts of uniquely mapped reads. To account for differences in sequencing depth between samples, we computed the expected breadth of coverage per species per sample using an established formula^8,19^. A species was considered present in a sample if it met both of the following criteria: (1) a ratio of observed to expected breadth of coverage exceeding 30% (considering that genomes were clustered at a species level using a 30% aligned fraction); and (2) an observed breadth of coverage >5% (to account for MAGs being up to 5% contaminated). These coverage thresholds and presence/absence criteria were previously validated using both mock community and synthetic datasets^19^. Species not meeting these thresholds were assigned a read count of zero.

To correct for potential study-specific batch effects, we applied a conditional quantile normalization algorithm implemented in ConQuR (v.1.2.0)^52^. Each of the 62 studies was evaluated as a potential reference for batch correction. Based on PERMANOVA results, study SRP188507 was selected as the final reference, as it yielded the lowest study effect size (R² = 0.088).

We assessed the association between microbiome alpha diversity and each disease condition using a linear mixed-effects model. The Shannon diversity index (rank-transformed) was used as the response variable, while disease status, sequencing depth and study source were used as fixed effects. If applicable, the subject ID was also used as a random intercept to account for repeated measures. The *P* value corresponding to the effect of disease status was extracted from the model summary and corrected for multiple testing across diseases using the Benjamini-Hochberg method.

### Machine learning modelling for disease classification

We developed supervised machine learning classifiers to assess the predictive potential of the microbiome for disease classification. For each disease, we conducted three types of analyses: (1) pooled analysis of studies from the same disease, (2) cross-study validation within the same disease, and (3) pairwise cross-disease validation (i.e., training the model with one disease and testing it on another). All machine learning analyses were performed using a custom workflow (https://github.com/alexmsalmeida/ml-microbiome) derived from the Mikropml R package^53^. For the pooled analysis, we evaluated three supervised learning algorithms — ridge regression, random forest, and gradient boosting — and selected ridge regression for all subsequent analyses based on the superior performance observed. Batch-corrected abundances of filtered microbiome species were transformed using the centred log-ratio (CLR) and used as input features for the machine learning models. Prior to modelling, features were filtered to exclude those with <1% prevalence across all samples or with zero variance. Model training and hyperparameter tuning were performed on 80% of the data using 5-fold cross-validation, while the remaining 20% was held out for testing using the best hyperparameters. This procedure was repeated 20 times with different random seeds to ensure robustness. To account for repeated measures in longitudinal datasets, subject IDs were used as a grouping factor to ensure that samples from the same individual were assigned exclusively to either the training or testing set. Model performance was assessed using the Area Under the Receiver Operating Characteristic Curve (AUROC).

### Differential species abundance analysis

To identify microbial biomarkers associated with health and disease in case–control cohorts, we employed two complementary generalized linear modelling approaches: ALDEx2 (v1.32.0)^26^ and MaAsLin2 (v1.14.1)^27^. We performed the analysis separately for each disease, with health status, age group, continent, sequencing depth, and study source included as covariates in the models. For ALDEx2, we applied the ‘aldex.clr’ function to perform a centred log-ratio (CLR) transformation on the abundance data (read counts), followed by ‘aldex.glm’ to model differential abundance between health and disease. For MaAsLin2, read counts were normalized by genome length and sequencing depth using total sum scaling, then log-transformed prior to modelling. In both methods, species with a prevalence below 1% across samples were excluded to reduce sparsity. Multiple testing correction was performed using the Benjamini–Hochberg method, and microbial species were considered significant only if they passed a false discovery rate (FDR) threshold <5% in both ALDEx2 and MaAsLin2.

### Meta-analysis across disease cohorts

We summarized species-level results per disease by first averaging the effect sizes obtained with ALDEx2 and MAasLin2. Thereafter, effect sizes were summed across diseases and normalized by the number of diseases in which the species was found non-significant, resulting in a combined effect size that prioritized species showing consistent significance across multiple diseases. To identify the most promising uncultured taxa linked to health, we further calculated an uncultured health score for each bacterial genus present in the UHGG. This metric was based on the product of three factors: (i) the total number of health-associated species in the genus (log_10_-transformed); (ii) the combined effect sizes normalized by the number of species in the genus; and (iii) the proportion of uncultured species. In both (ii) and (iii), the denominator included only species within the genus that had a minimum prevalence of 1% in at least one case-control cohort.

To validate our approach using an alternative methodology, we performed an independent meta-analysis using the classical Stouffer’s method. A limitation of this approach is that it cannot accommodate the initial FDR thresholding applied when intersecting the ALDEx2 and MaAsLin2 results. Therefore, for this validation we relied solely on ALDEx2, which is recognized as the most conservative differential abundance method^54^. Using this approach, the ALDEx2 effect sizes obtained per species were weighted across diseases by the square root of the corresponding sample size, and Z-scores were derived from the reported *P* values. The weighted Z-scores were then aggregated to obtain a meta-analysis Z-score, which was converted into a combined *P* value. Statistical significance was assessed at an FDR threshold of 5%.

### Ecological modelling in healthy populations

To conduct a dedicated ecological analysis of the healthy human microbiome, we compiled all control samples from the case-control studies included in this work (*n* = 3,614), along with an additional dataset of 2,443 metagenomes from healthy subjects. We excluded all samples from infants and children under 13 years of age. Microbial correlation networks were constructed using FastSpar (v.1.0)^55^, a C++ implementation of SparCC^56^, based on species with a prevalence greater than 1%. Abundance values used in FastSpar were derived from the batch-corrected read counts. Co-abundance networks were generated separately for samples from five continents (Asia, Africa, Europe, North America and South America), with samples from Oceania excluded due to insufficient sample size. Thereafter, within each network we calculated exact *P* values from 1,000 bootstrap replicates, applying a Benjamini-Hochberg multiple testing correction to retain correlations with FDR <5%. SparCC correlation values were transformed into distances (calculated as 1 − |correlation|), and for each species, we computed three network centrality measures — betweenness, closeness, and degree — using the igraph R package^57^. Betweenness centrality reflects how often a species lies on the shortest paths between others, closeness centrality indicates how close a species is to all others in the network, and degree centrality represents the number of direct connections a species has. Previous studies have shown that species with high closeness, high degree and low betweenness are more likely to be keystone^28,29^. Therefore, we rescaled each metric with betweenness values inverted (multiplied by −1), and averaged across continents to derive a single score, referred to as keystonness. Species within the top 1% keystonness score were classified as ‘keystone’. Results were repeated by further filtering the initial co-abundance networks to only retain edges with |correlation| > 0.1 and with |correlation| > 0.2.

### Functional prediction of candidate biomarkers

Functional prediction analyses were performed on the candidate biomarkers of health and disease using genome-based annotation methods. For this analysis, we excluded species that exhibited a conflicting signal (that is, linked to health in some disease cohorts and to disease in others). Protein-coding sequences were first extracted from the genome assemblies using Prokka (v1.14.16)^58^. Each genome was then functionally characterized using eggNOG-mapper (v2.1.3)^59^ and KOFams (release 2021-11)^30^ with default parameters. A generalized linear model was applied to identify KEGG Orthologs (KOs) significantly associated with health- or disease-related biomarkers (using the ‘glm’ function in R, family = "binomial"), while controlling for taxonomy (Order level) and genome type (MAG or isolate) as covariates. Exact *P* values were corrected for multiple testing using the Benjamini-Hochberg method and filtered based on an FDR <5%.

### Oscillospiraceae functional comparison

A phylogenetic and functional analysis of the Oscillospiraceae family was performed to identify features specifically associated with the genus CAG-170. First, a family-level phylogenetic tree was constructed using all high-quality (completeness >90%) species genomes of Oscillospiraceae present in the UHGG collection (*n* = 152). The tree was generated with FastTree (v.1.2.11)^60^ based on the alignment of bacterial marker genes obtained using GTDB-Tk (v.2.3.2, database release 214)^50^.

Functional annotation was performed for all high-quality (>90% complete), non-redundant (99.9% average nucleotide identity) Oscillospiraceae genomes (*n* = 4,025), following the same approach used for characterisation of the species biomarkers. To identify functional traits associated with the CAG-170 genus, a mixed-effects logistic regression analysis was carried out for each KEGG Ortholog (KO) predicted across these genomes. Only KOs found at a prevalence >1% were included. In fitting the model, the response variable was defined as the presence or absence of the KO in each genome, and the predictor variable was genus membership (CAG-170 vs. other Oscillospiraceae). We included as a random effect the species affiliation to account for the fact there were multiple genomes from the same species under analysis. *P* values from all models were corrected for multiple testing and KOs with an FDR <5% were considered statistically significant.

### Characterization of CAG-170 in a longitudinal IBD cohort

We analysed 1,118 metagenomic samples from a longitudinal IBD cohort of the Human Microbiome Project (HMP2-IBD)^9^ to further investigate the abundance and subspecies genetic diversity of the CAG-170 taxon over time in health and disease. This cohort included individuals diagnosed with Crohn’s disease, ulcerative colitis, and non-IBD controls. Participants were originally followed for one year, with stool samples collected approximately every two weeks. Each microbiome sample was associated with a ‘dysbiosis’ score, quantifying the deviation from a healthy gut microbial composition as defined in the original HMP2 study.

To evaluate the association between CAG-170 abundance and ‘dysbiosis’, we applied a linear mixed-effects model. Specifically, we modelled the CAG-170 genus-level abundance as a function of the dysbiosis score, with subject ID as a random effect to account for repeated measurements within subjects. The significance of the association between dysbiosis and CAG-170 abundance was assessed using an ANOVA test on the fitted linear mixed-effects model. The analysis was then repeated by partitioning dysbiosis scores into ‘high’ or ‘low’ based on the median value.

We further investigated the subspecies genetic diversity of CAG-170 within this cohort. First, we extracted all accessory genes from the Unified Human Gastrointestinal Protein (UHGP-90-HQ)^18^ catalog exclusive to each of the 13 CAG-170 species, with accessory genes being defined as those present in fewer than 90% of conspecific genomes. For this purpose, only genomes with >90% completeness and <5% contamination, as defined in the UHGG database, were considered. To screen for the presence of these accessory genes in the HMP2-IBD study, metagenome assemblies were generated for all 1,118 HMP2-IBD metagenomic samples using MEGAHIT (v1.2.9)^61^ with the ‘--min-contig-len 500’ option. Thereafter, protein-coding sequences were predicted from these assemblies using Prodigal (v2.6.3)^62^ with the ‘-p meta’ parameter. To profile the gene presence across samples, we used DIAMOND (v2.1.8)^63^ in ‘blastp’ mode to align all extracted CAG-170 accessory genes against the predicted protein sequences from the HMP2-IBD assemblies. Alignments were filtered using a threshold of ≥90% amino acid identity and ≥80% coverage of the shorter sequence. This resulted in a binary presence/absence matrix of accessory genes across all metagenomic samples. We kept only accessory genes that appeared in more than 1% of HMP2-IBD samples, and retained only those samples that contained at least one of these genes for downstream analysis. To assess the association between gene presence and dysbiosis, a mixed-effects logistic regression model was fitted, with subject ID as a random effect. *P* values were adjusted for multiple testing using the Benjamini–Hochberg method, and genes with FDR <5% were considered statistically significant. Functional annotations, including eggNOG and Clusters of Orthologous Groups (COG)^64^ categories for the candidate accessory genes were obtained from the original UHGP catalog.

## Data availability

The human gut metagenomic samples used in this study are publicly available in the European Nucleotide Archive (see Supplementary Table 1 for all associated accession codes). The sequence databases used were originally retrieved from the Unified Human Gastrointestinal Genome (UHGG) catalog v.1.0 and the Unified Human Gastrointestinal Protein (UHGP-90-HQ) catalog v.1.0. Abundance data estimated for the UHGG species and all metagenomic samples here included are available in figshare at https://doi.org/10.6084/m9.figshare.29963597.v1. The protein FASTA file of the CAG-170 accessory genes detected in the HMP2-IBD study can be accessed from https://doi.org/10.6084/m9.figshare.29963588.v1. Accession code of the human reference genome used for decontamination (GRCh38) is GCA_000001405.15.

## Code availability

Custom scripts and pipelines used in this work are publicly available in GitHub at https://github.com/microfundiv-lab/SecretBugs.

## Extended Data Figures

**Extended Data Figure 1.**
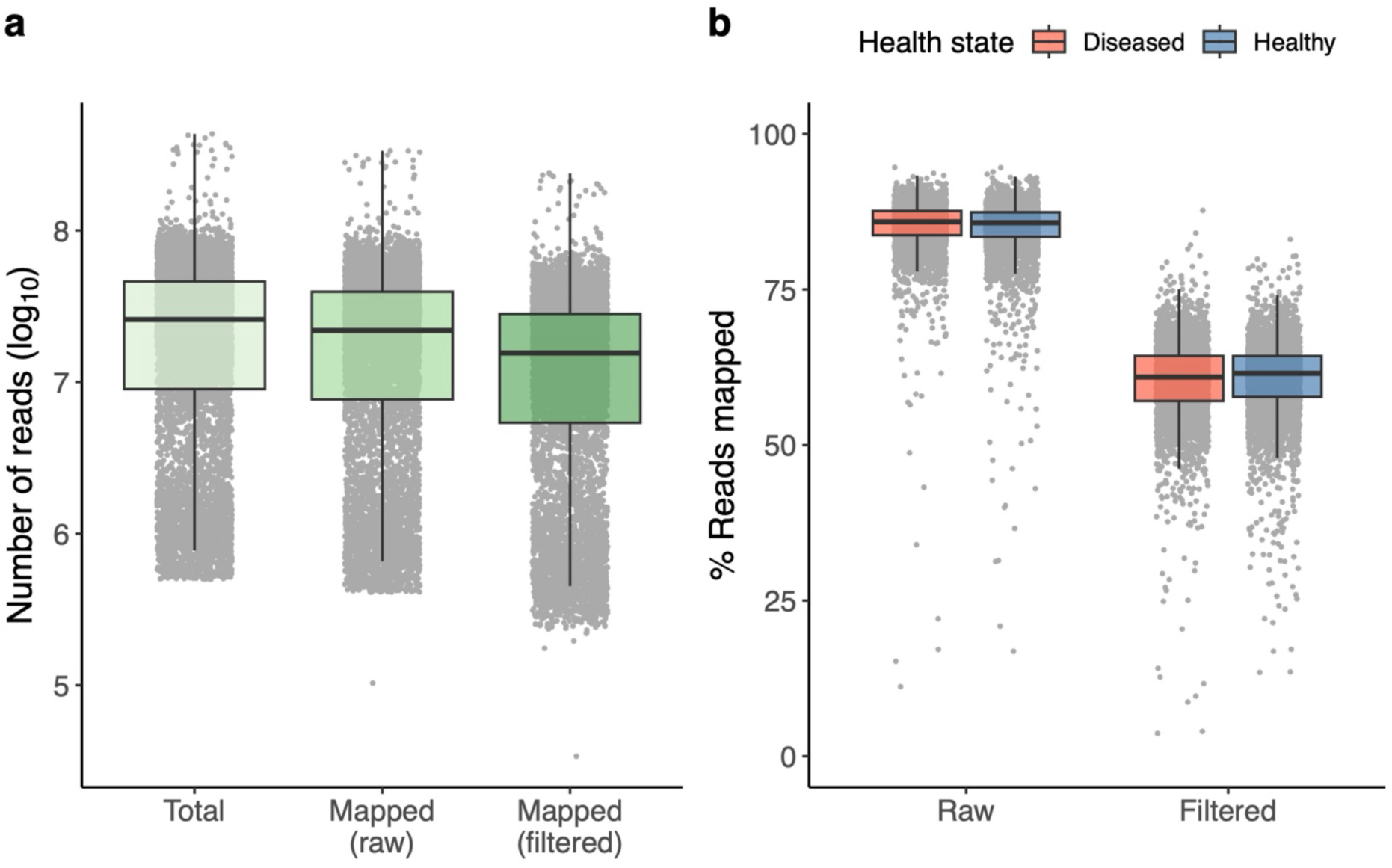
Distribution of sequencing depth and mapping rates. **a,** Distribution of the total number of reads, number of reads mapped (raw) and number of reads mapped after filtering among the case-control metagenomic samples (*n* = 8,672). **b**, Proportion of reads mapped across healthy and diseased samples without applying any mapping filters (“Raw”) and after applying filters based on observed and expected breadth of coverage (“Filtered”). The centre line within the box represents the median score. Whiskers are shown extending to the furthest point within 1.5 times the IQR from the box (disease samples, *n* = 4314; healthy samples, *n* = 4358).

**Extended Data Figure 2.**
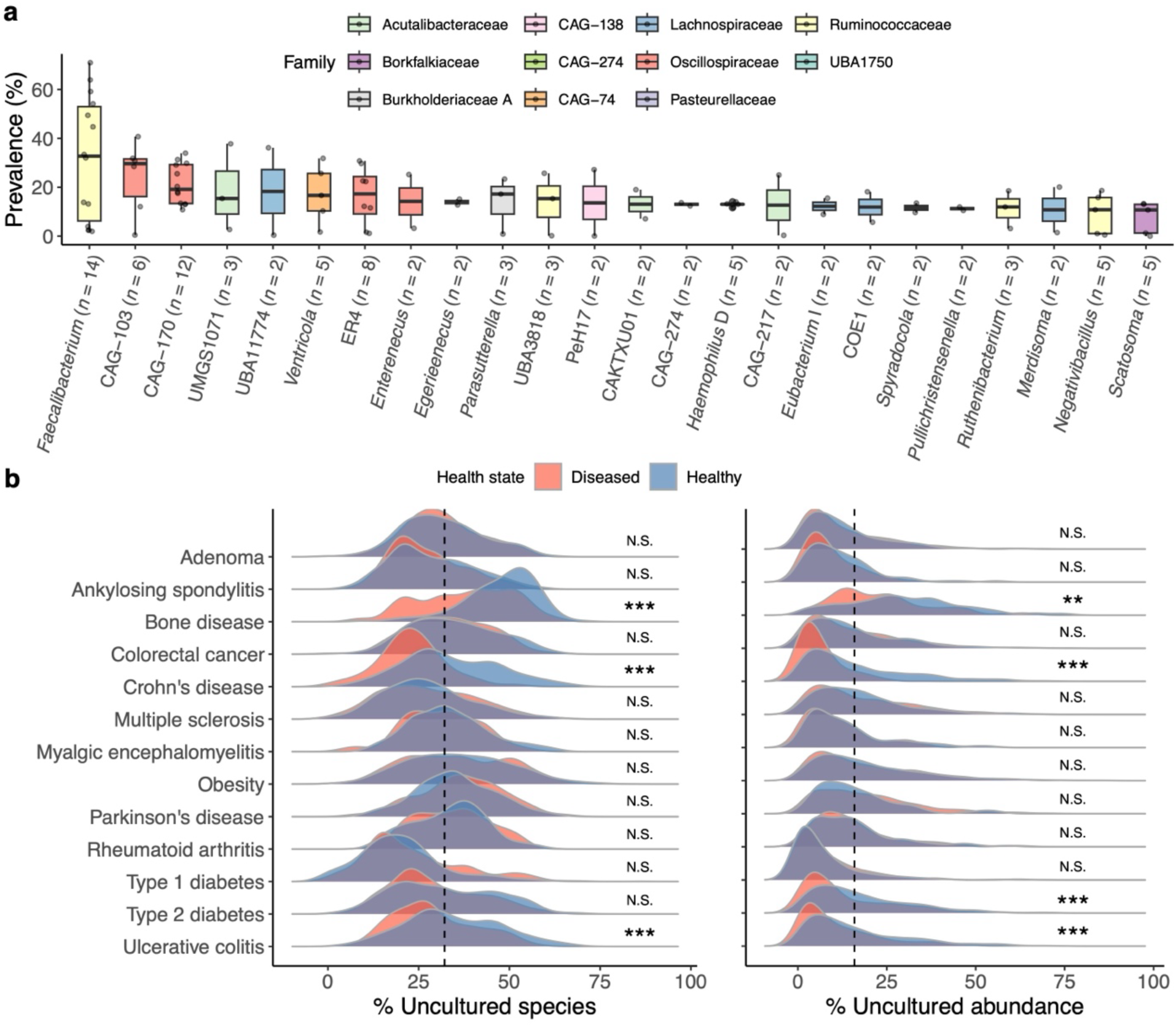
Profiling uncultured species across multiple disease conditions. **a,** Prevalence of uncultured species, partitioned by genus, shown for the 50 most prevalent genera. Taxa labels are in accordance with the Genome Taxonomy Database (GTDB) release 214. The centre line within each boxplot represents the median score. Whiskers are shown extending to the furthest point within 1.5 times the IQR from the box. Number of data points (species) is labelled next to each genus. **b**, Proportion of uncultured species (left) and uncultured abundance (right) detected across all case-control studies. The proportion of uncultured species was calculated as the number of uncultured species detected out of all species present in the sample, whereas uncultured abundance was inferred by dividing the number of reads mapped to uncultured species out of all reads mapped to the UHGG. Vertical dashed lines represent the median values. *P* values were derived from a two-sided Wilcoxon rank-sum test and corrected for multiple testing using the Benjamini-Hochberg method. **FDR < 0.01; ***FDR <0.001; N.S. = Non-significant.

**Extended Data Figure 3.**
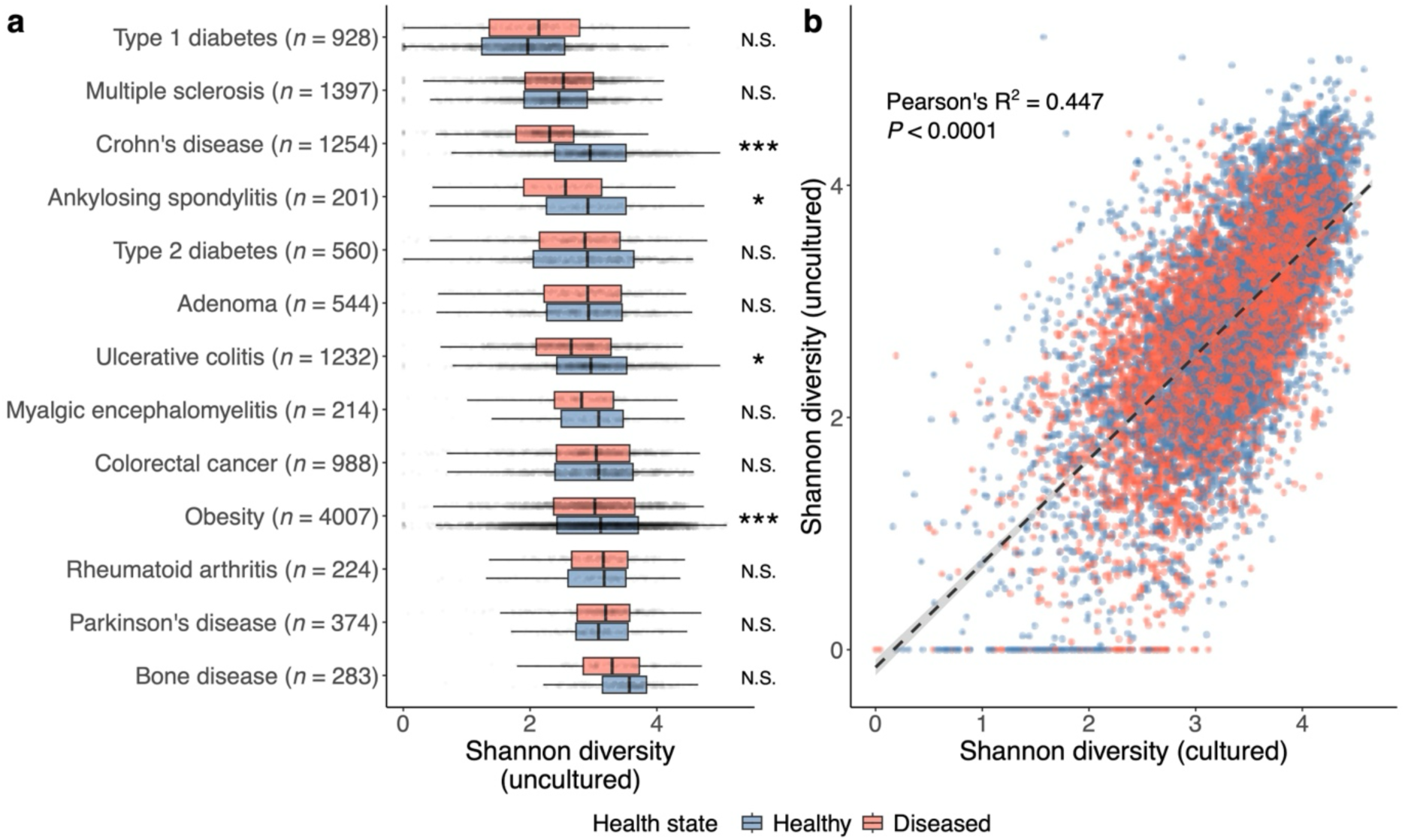
Alpha diversity estimates in health and disease. **a**, Distribution of alpha diversity values (Shannon index) estimated using only uncultured species across all the diseases tested. *P* values were derived from a linear mixed-effects model with Shannon diversity rank as the response, and health state, study source, and read count (log-transformed) as fixed effects. The subject ID was used as a random intercept if there was more than one sample per subject. *adjusted *P* < 0.05; ***adjusted *P* < 0.001; N.S. = Non-significant. The centre line within the box represents the median score. Whiskers are shown extending to the furthest point within 1.5 times the IQR from the box. Total number of samples (*n*) per disease are indicated in parenthesis. **b,** Correlation between the Shannon alpha diversity estimated using either cultured or uncultured species. Samples are coloured by health state.

**Extended Data Figure 4.**
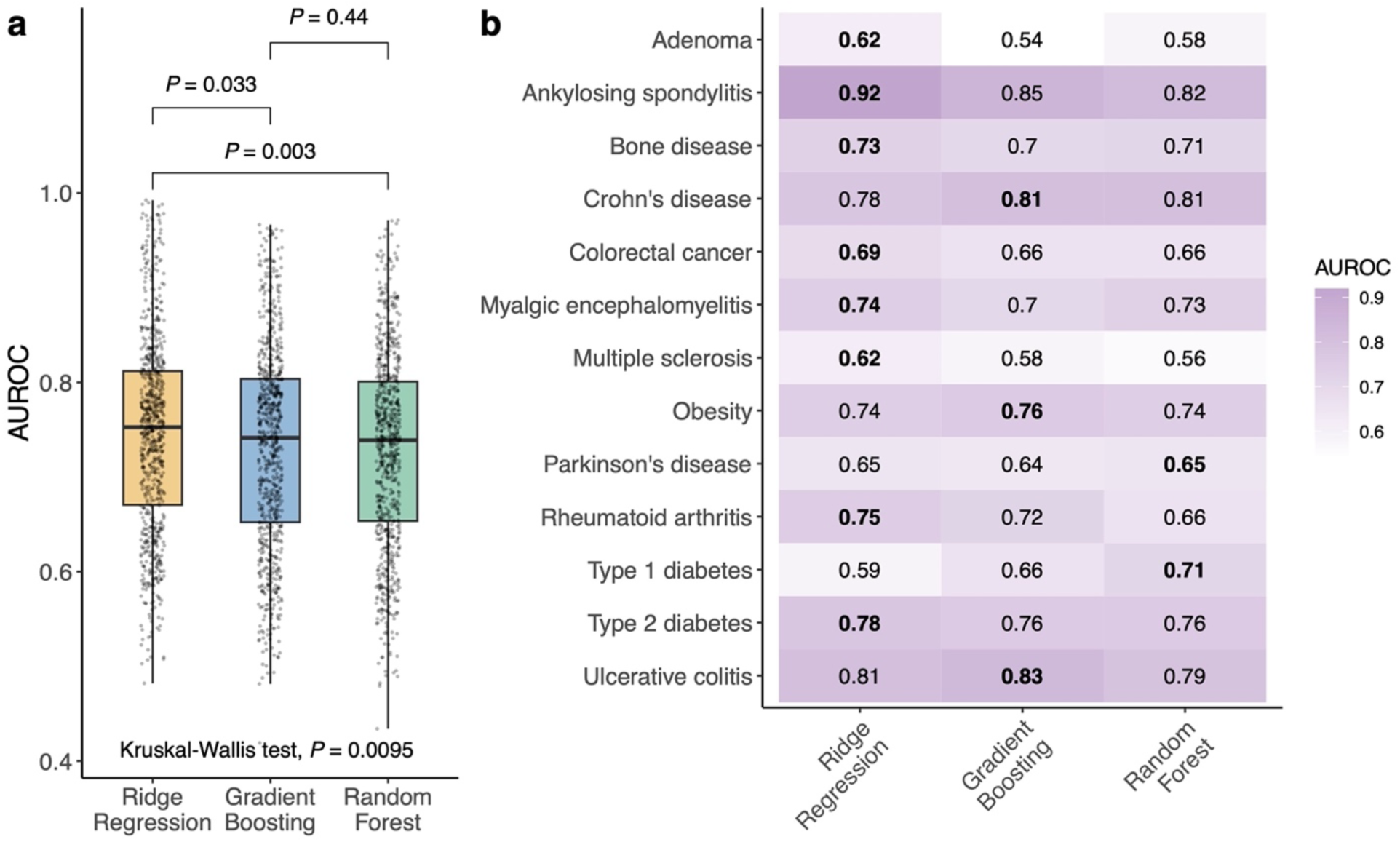
Evaluating the best supervised machine learning model. **a,** Model performance assessed by the Area Under the Receiver Operating Characteristic Curve (AUROC) for the three supervised machine learning methods tested. The centre line within the box represents the median score (*n* = 780, corresponding to 20 seeds per model per dataset — cultured, uncultured and both — across 13 diseases). Whiskers are shown extending to the furthest point within 1.5 times the IQR from the box. *P* values were derived from a two-sided Wilcoxon rank-sum test and corrected for multiple testing using the Bonferroni–Holm method. **b**, Performance of the models when using the uncultured microbiome alone, partitioned by disease. Values correspond to the median value across 20 seeds. The best performing model per disease is indicated in bold.

**Extended Data Figure 5.**
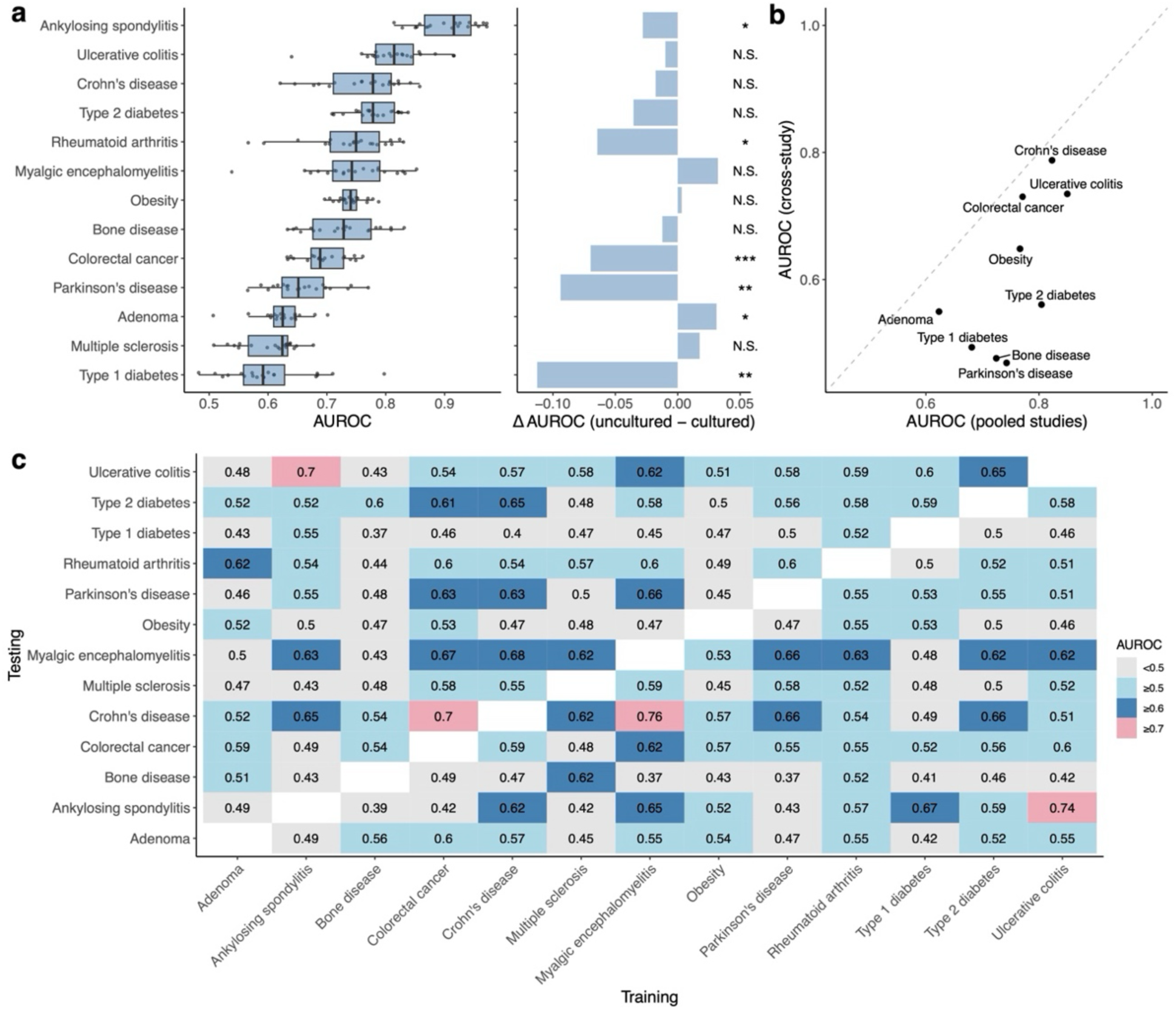
Disease classification using machine learning. **a**, Performance of the machine learning models using uncultured species as features (left) and compared to those using only cultured species (right). The centre line within the box represents the median score (*n* = 20 seeds per model). Whiskers are shown extending to the furthest point within 1.5 times the IQR from the box. On the right, bars with positive values indicate that models based on uncultured species outperformed those based on cultured species only, and vice-versa. *P* values were derived from a two-sided Wilcoxon rank-sum test and corrected for multiple testing using the Benjamini-Hochberg method. *FDR < 0.05; **FDR < 0.01; ***FDR <0.001; N.S. = Non-significant. **b**, Comparison of model performance (median AUROC) when pooling studies from the same disease versus a cross-study validation (training on one set of studies and testing on others). Diagonal dashed line represents equal performance between both approaches. **c**, Model performance of a cross-disease validation, where models are trained on one disease and tested on another. AUROC values shown are medians across 20 random seeds.

**Extended Data Figure 6.**
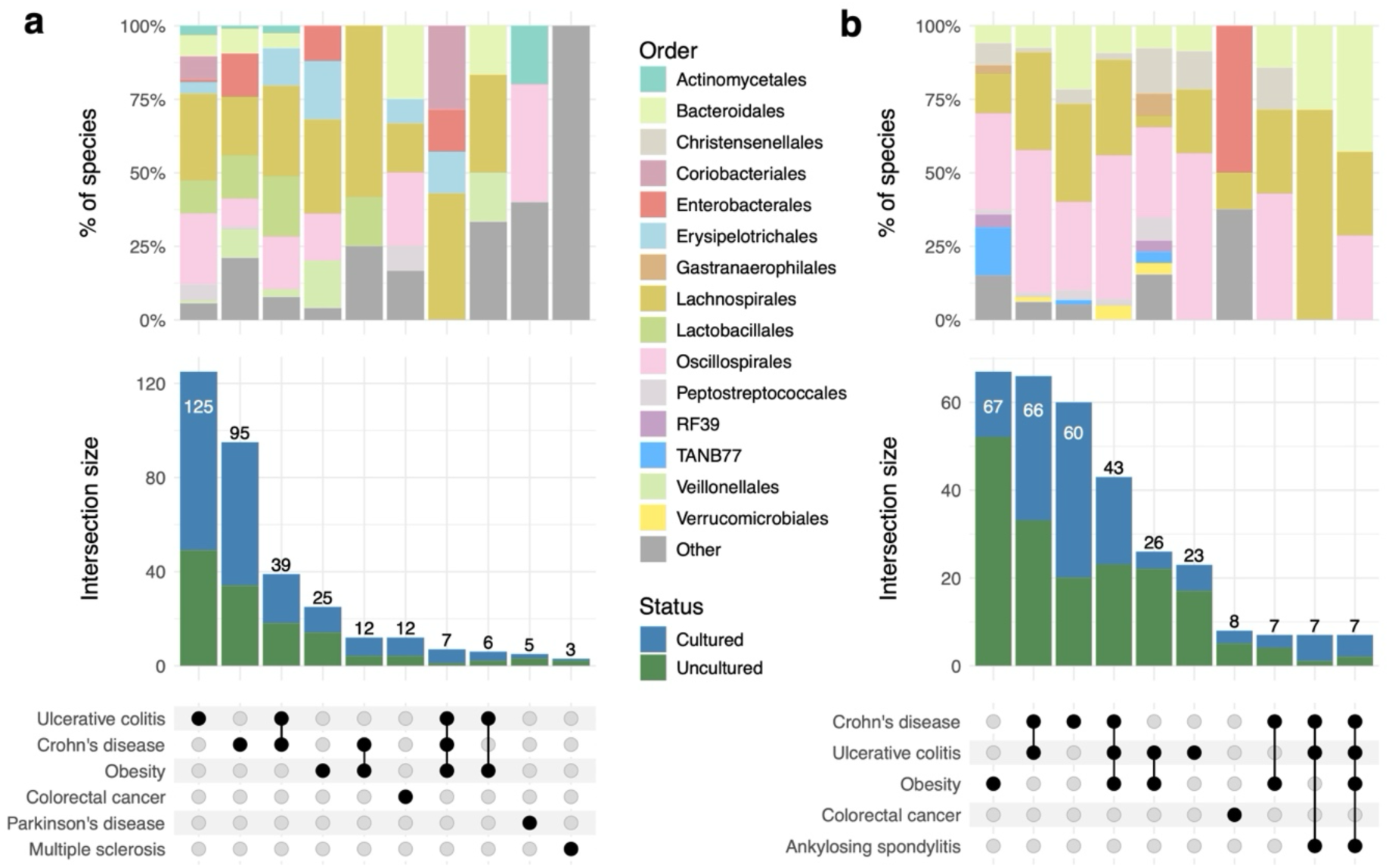
Overlap of health and disease biomarkers across multiple conditions. **a**, Number of prokaryotic species classified as disease biomarkers (higher abundance in disease) and the level of overlap across multiple conditions. Vertical bars represent the number of biomarkers found in the diseases highlighted with black dots in the lower panel (only the top 10 intersections are shown). Stacked bar plots at the top display the taxonomic affiliation (order rank) of the species within each intersection. **b**, Same representation as panel (**a)** but only considering health biomarkers (higher abundance in healthy controls).

**Extended Data Figure 7.**
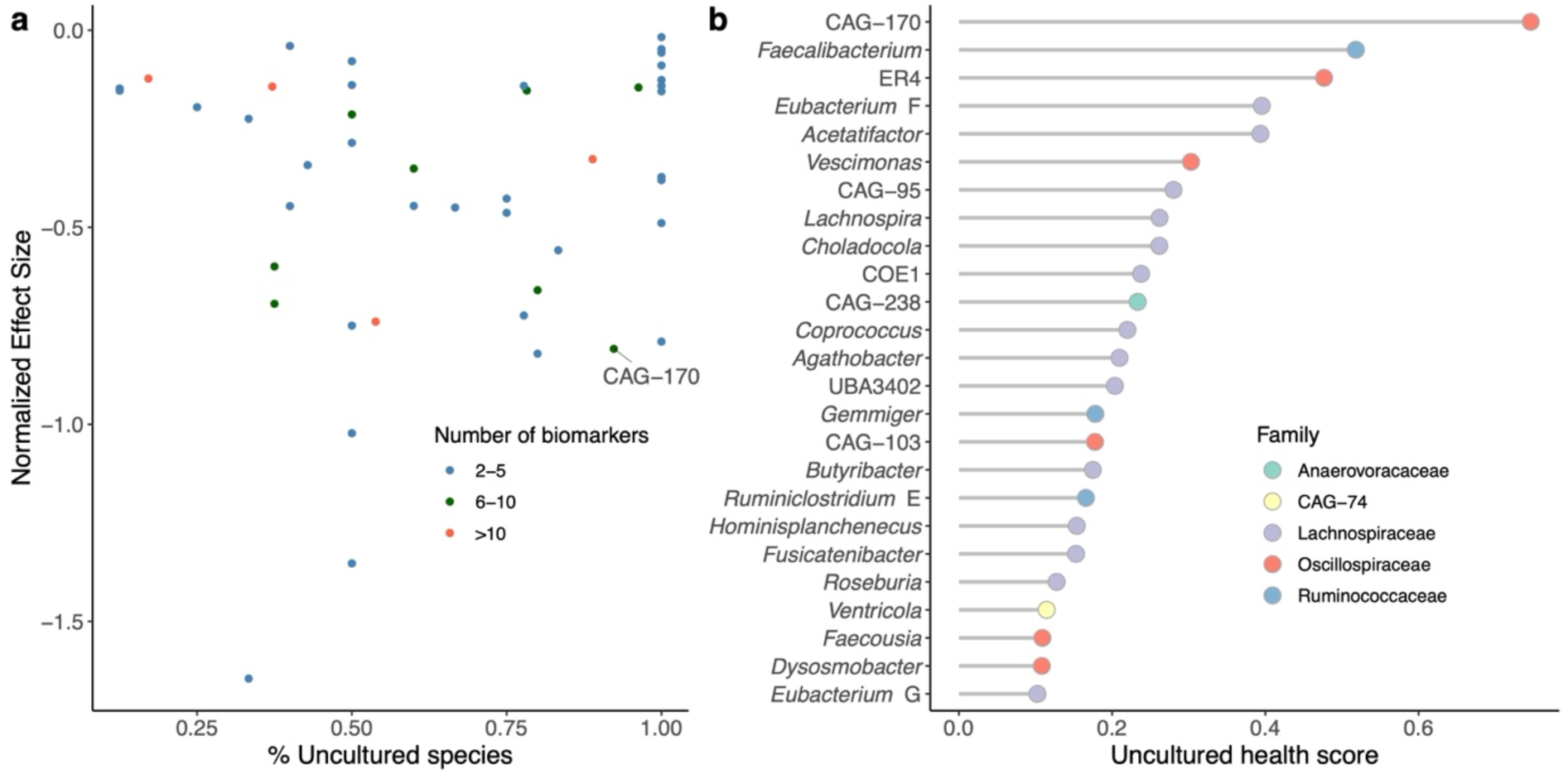
Validation of uncultured health scores using the Stouffer’s method. **a,** Proportion of uncultured species relative to the estimated effect size for each bacterial genus in the UHGG. A negative effect size indicates an association with health. Each genus is coloured according to the number of species significantly associated with health. Effect sizes and *P* values for defining statistical significance were combined across diseases using the Stouffer’s method. **b**, Bacterial genera ranked by the absolute value of their uncultured health score, calculated as the product of the number of significant species (log_10_-transformed), normalized effect size, and proportion of uncultured species.

**Extended Data Figure 8.**
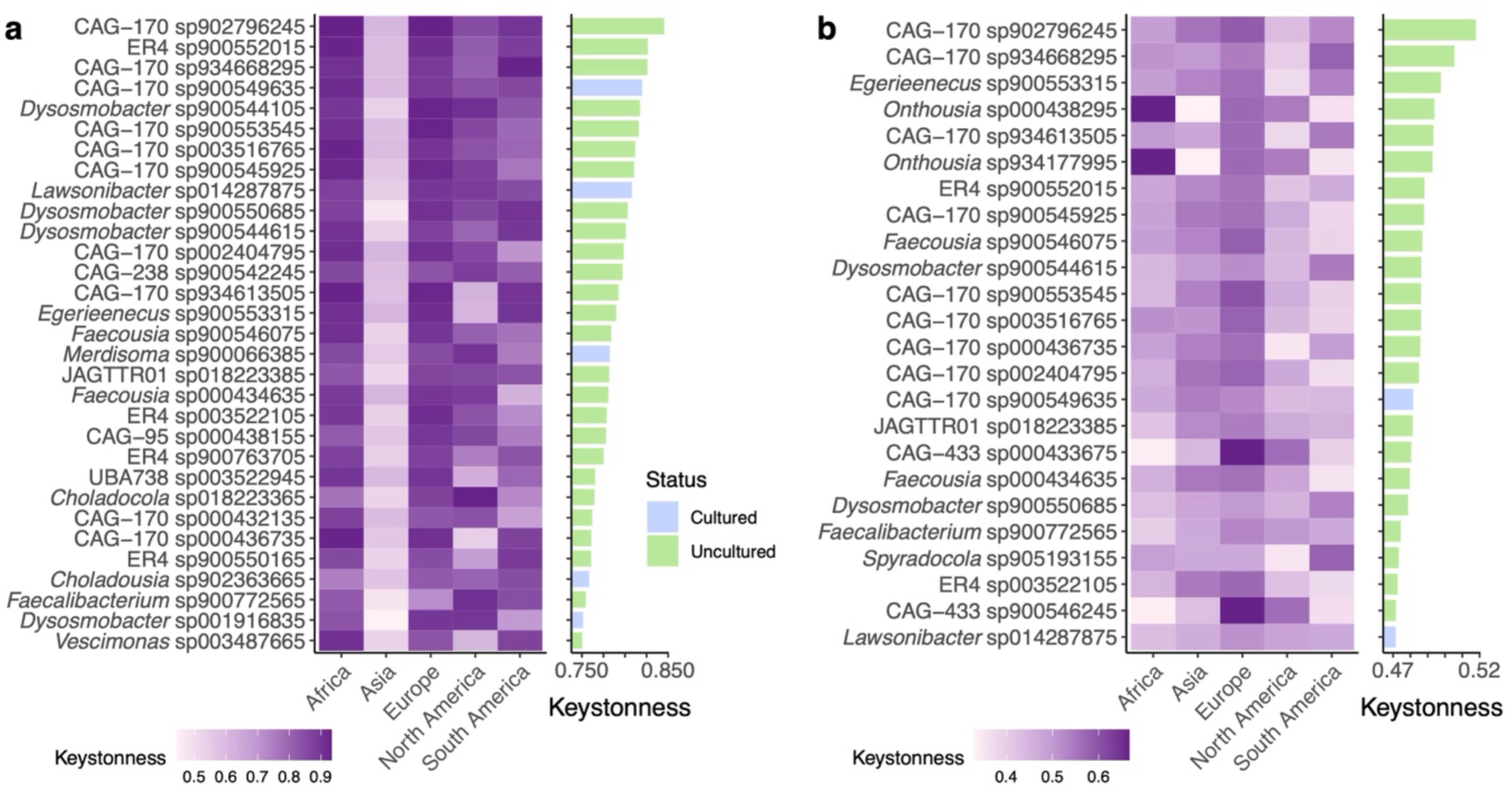
Comparison of keystone species following additional correlation filters. **a,** Keystonness scores of the top 1% most central species (defined as ‘keystone’) after filtering network edges with a minimum |correlation| of 0.1. The heatmap shows continent-specific values, and the bar plot shows the mean keystonness across continents. **b**, Same representation as panel (**a**) but with networks filtered with a minimum |correlation| of 0.2. The genus CAG-170 is the most represented taxon within both sets.

**Extended Data Figure 9.**
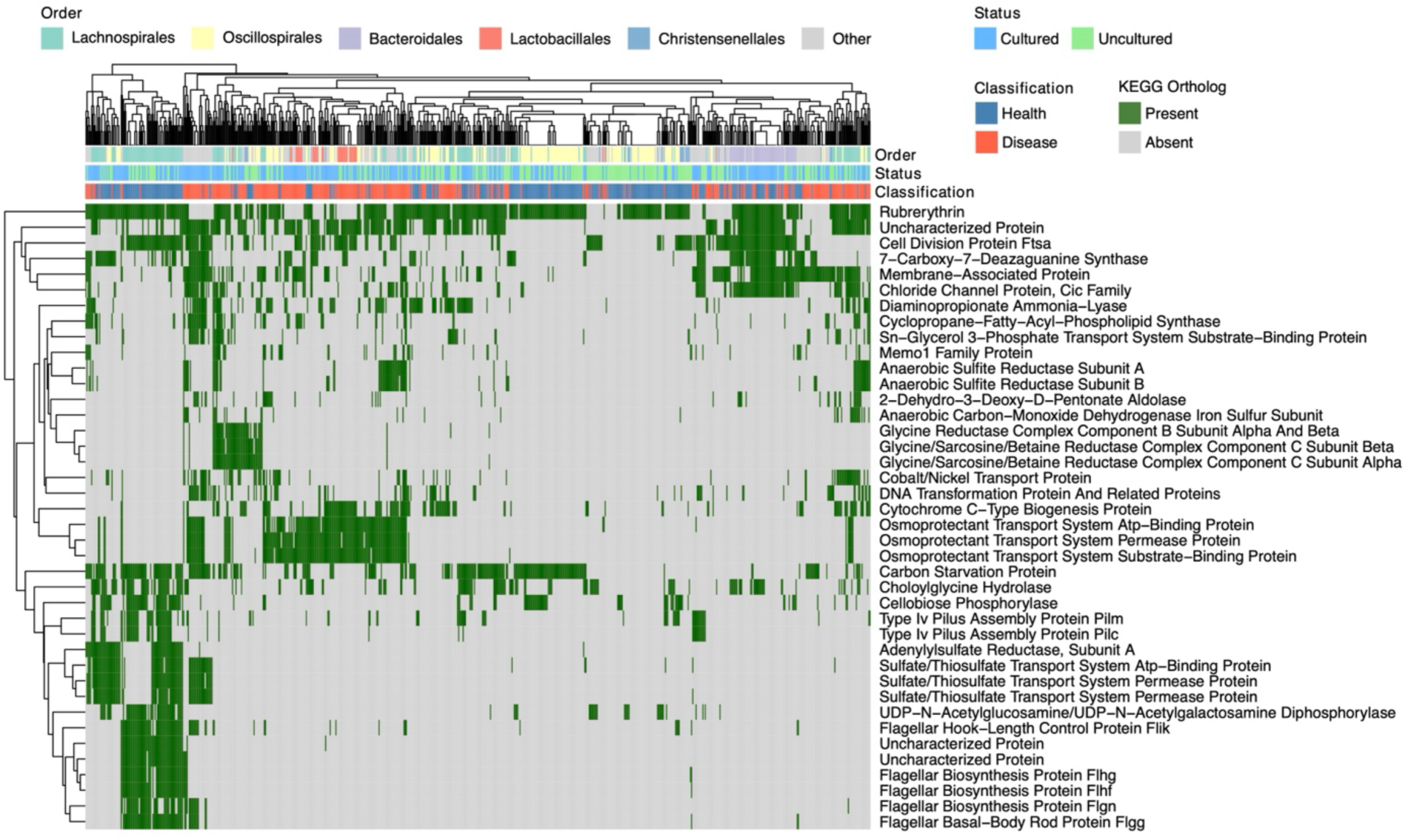
Functional signatures of health and disease biomarkers. Heatmap depicting the distribution of the top 20 KEGG Orthologs (KOs) associated with health and disease biomarkers. Columns represent bacterial species coloured by their taxonomic affiliation, genome type and classification (health or disease). KOs and genomes are clustered using a complete linkage hierarchical clustering on the basis of their presence/absence patterns.

## Notes

https://doi.org/10.6084/m9.figshare.29963597.v1

https://doi.org/10.6084/m9.figshare.29963588.v1

https://github.com/microfundiv-lab/SecretBugs

### Competing Interest Statement

The authors have declared no competing interest.

